# Kilohertz electrical stimulation drives widespread neuronal activation and temporal interference imposes focal rhythmic modulation

**DOI:** 10.64898/2026.10.15.759497

**Authors:** Ryohei Yoshimoto, Matthew A. Stern, Thomas Eggers, Emma Acerbo, Nealen Laxpati, Ken Berglund, Claire-Anne Gutekunst

**Affiliations:** Department of Neurosurgery, Emory University School of Medicine, Atlanta, GA, United States; Coulter Department of Biomedical Engineering, Emory University and Georgia Institute of Technology, Atlanta, GA, United States; Children’s Healthcare of Atlanta, Atlanta GA

## Abstract

Temporal interference (TI) stimulation has attracted attention as a non-invasive approach that can target deep brain structures through interacting kilohertz-frequency electric fields. Its basic premise is that TI stimulation evokes neuronal responses through low-frequency amplitude modulation (AM), while the underlying kilohertz-frequency stimulation is ineffective to induce activity, the latter of which has not been tested rigorously at the cellular level. Here, we combined two-photon imaging in mouse models with immunohistochemistry and computational modeling to characterize spatiotemporal patterns of neuronal responses during TI stimulation. We found that kilohertz-frequency stimulation was sufficient to evoke activity without AM. In contrast, AM organized neuronal activity into rhythmic responses at AM frequency, with the strongest entrainment near the center of interference. Thus, kilohertz-frequency stimulation can drive widespread neuronal activation, upon which TI imposes focal rhythmic modulation. Our results provide a framework for refining TI stimulation and highlight the potential of kilohertz-frequency stimulation for brain stimulation.

## Introduction

Neuromodulation has become an important approach in basic neuroscience and clinical therapy because it enables direct manipulation of neural activity for both probing function and treating neurological and psychiatric disorders. Among neuromodulatory techniques, deep brain stimulation (DBS) has shown strong therapeutic potential by directly targeting deep brain structures and modulating pathological neural circuits^1,2^. However, because DBS requires surgical implantation of electrodes into the brain, this technique is inherently invasive and carries risks associated with surgery, limiting its accessibility and acceptance amongst patients^3,4^. In contrast, non-invasive brain stimulation techniques, such as transcranial direct current stimulation (tDCS), transcranial alternating current stimulation (tACS), and transcranial magnetic stimulation (TMS), allow modulation of brain activity without surgical intervention^5–7^. These methods are attractive because they are relatively safe and accessible. However, a major limitation of conventional non-invasive stimulation is its limited ability to selectively reach deep brain targets^8,9^. Therefore, there is a growing need for neuromodulatory approaches that can target deep brain structures while preserving the advantages of non-invasiveness.

To overcome these limitations, temporal interference (TI) stimulation has been proposed as a promising approach for non-invasive deep brain stimulation. The theoretical framework of TI is based on the interaction of two kilohertz high-frequency electric currents delivered through two pairs of electrodes at offset frequencies. Their intersection within the tissue generates an amplitude-modulated (AM) electric field at depth, with an envelope that oscillates at the frequency difference between the two currents. By adjusting the spatial distribution and relative strength of the two fields, the region of strongest AM electric field can be positioned within a desired deep brain target, while the surrounding tissues are exposed predominantly to the individual kilohertz carrier fields. This approach assumes that neurons respond primarily to the low-frequency envelope, but not to the unmodulated kilohertz carrier signals as they are unable to fire action potentials at such a high frequency^10^. This selective sensitivity has been attributed to the low-pass filtering properties of the neuronal membrane, which are thought to attenuate responses to high-frequency inputs while preserving sensitivity to slower modulation components^11^. As a result, TI was expected to selectively activate target regions where AM is strongest, while reducing off-target activation in surrounding areas near the stimulation electrodes, where the low frequency envelope is less formed. Because of this potential to target deep brain regions, TI stimulation has drawn considerable attention in the neuromodulation field^12^. Research has rapidly expanded from rodent^10^ and non-human primate models^13,14^ to human studies targeting deep brain regions, such as the hippocampus^15,16^ and striatum^17–19^. Early clinical investigations have also begun to explore its therapeutic potential in treating neurological and neuropsychiatric disorders^12,20–23^.

Despite the considerable attention TI stimulation has received, several fundamental questions remain unresolved. One of the most important unresolved characteristics is the spatiotemporal characteristics of neural responses to TI stimulation. In terms of spatial focality, it remains unclear whether actual neural activation truly corresponds to the computationally predicted distribution of the AM. Previous studies have inferred spatial patterns of neural activation from AM maps derived from computational or experimental field measurements, rather than directly measuring neuronal activity^14,15,17,18^. Conventional approaches for whole-brain imaging, such as fMRI, lack single-neuron resolution and therefore cannot capture fine spatial patterns of neural activity during TI stimulation^15–18,24,25^. *Ex vivo* studies using patch-clamp recordings in brain slice preparations provide single-cell resolution, although they are low throughput and do not adequately address the spatial gradient of AM across the field ^26–30^. As a result, neural responses to highly variable AM remain largely unexplored. In addition, it is not yet clear whether neurons are rhythmically synchronized to the envelope of TI stimulation at the single-neuron level. Although TI is expected to modulate neural activity at AM frequency, direct evidence of this at the individual neuron level remains limited. Furthermore, it is uncertain whether such rhythmic activity occurs selectively near the predicted interference region or is more broadly distributed across a stimulated tissue volume.

The seminal publication on TI stimulation reported that neuronal responses were insignificant when stimulated with kilohertz carriers alone and probed through intracellular recordings of patch-clamp electrophysiology and expression of an immediate early gene histologically^10^. However, since the time of Bernard Katz, kilohertz sinewaves have been used to excite nerves^31^, and, in fact, modern computational and experimental studies suggest that neurons can respond to kilohertz-frequency stimulation in the brain as well as in the peripheral nerves^26,27,32–37^. In addition to neural activation, high-frequency fields are known to produce conduction block at high intensities^38^. Thus, it seems important that the effect of weakly modulated or unmodulated stimulation outside the predicted TI focus needs to be carefully reassessed using a high-throughput functional assay.

While studies of TI stimulation in animal models – whether in brain slice preparation or *in vivo* – primarily rely on readouts of neuronal spiking, such suprathreshold stimulation is considered implausible in human conditions^30,35,39–41^. Thus, in this study, we employed methodologies to measure both suprathreshold spiking to high-amplitude TI stimulation as well as subthreshold fluctuation of membrane potential to weaker stimuli, the latter of which better reflect actual use.

To characterize spatiotemporal properties, neural responses to high-frequency stimulation, and subthreshold effects of TI, we combined functional optical imaging approaches *ex vivo* and *in vivo* in a mouse model supplemented with *in silico* simulations and immunohistology. We used two-photon imaging with both genetically encoded calcium and voltage sensors which allow for simultaneously monitoring individual cell responses while avoiding interference from electrical stimulation artifacts. We found that unmodulated high-frequency stimulation alone was sufficient to evoke neural activity, while AM primarily organized that activity in time. Although neuronal activation was broadly distributed across the imaging field, beat-frequency entrainment remained spatially localized near the interference center. Furthermore, TI induced rhythmic depolarization of the membrane potential, demonstrating temporal modulation at the subthreshold levels. We hope that our study will provide a useful framework when designing and implementing TI stimulation in animals and humans and motivate further exploration of kilohertz-frequency electric fields as a modality for brain stimulation.

## Result

### High-frequency stimulation alone is sufficient to induce neuronal responses independent of amplitude modulation

TI is generated by the interaction of two high-frequency electric fields with slightly different frequencies, producing an AM waveform whose envelope fluctuates at the frequency difference. We defined the strength of this envelope modulation as an AM index using the following calculation: (Env_max_ – Env_min_) / Env_max_ × 100%, where Env_max_ and Env_min_ are the maximum and minimum of the amplitude of the envelope, respectively. AM index and amplitude during TI stimulation continuously vary across space. In this study, in addition to TI electric fields generated by two pairs of electrodes, we also used a simplified stimulation generated by one pair of electrodes that mimics the waveform that would develop in the target structure with predefined AM index and amplitude (premodulated TI or pmTI)^42^. In pmTI stimulation, amplitude-modulated waveforms with specified AM indices are generated in advance, enabling us to examine how local waveform features shape neuronal responses across the space (Fig. 1a).

**Fig. 1.**
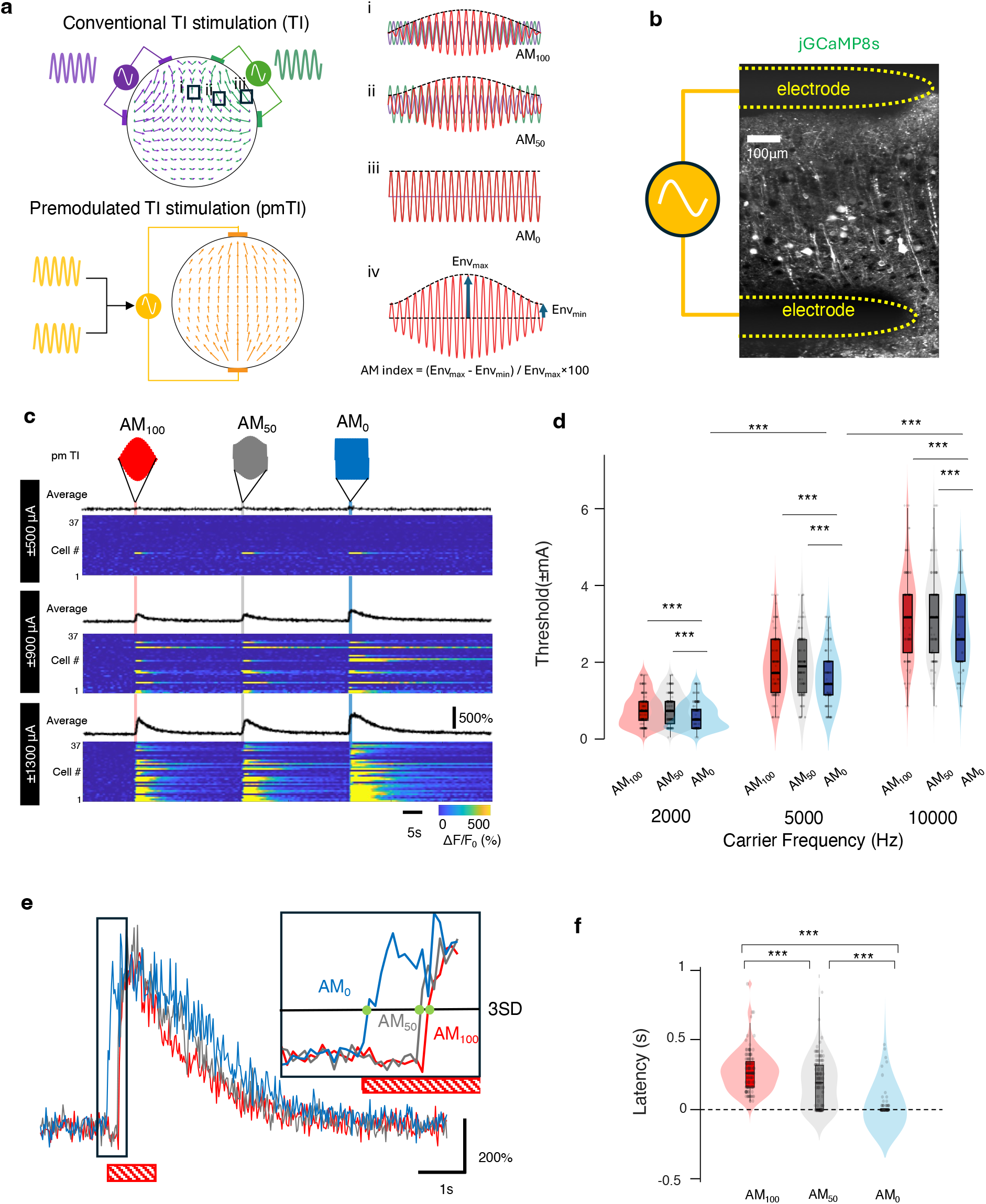
Neuronal responses during high-frequency stimulation with or without amplitude modulation. **a**, Conceptual comparison of conventional temporal interference (TI) stimulation (top left) and premodulated TI (pmTI) stimulation (bottom left). In conventional TI stimulation, two high-frequency sinusoidal electric fields with slightly different carrier frequencies spatially interact, producing location-dependent amplitude modulation. i–iii, Representative waveforms at different spatial locations within the TI field, corresponding to AM indices of 100%, 50%, and 0%, respectively. In pmTI stimulation, two sinusoidal waveforms are summed before delivery to generate a single waveform with a predefined AM index, producing the same waveform throughout the stimulation field. iv, Representative waveform at an arbitrary location within the pmTI field, illustrating the definition of the AM index. The AM index was calculated as (Env_max_ − Env_min_) / Env_max_ × 100, where Env_max_ and Env_min_ represent the maximum and minimum envelope amplitudes, b, Experimental configuration for pmTI stimulation using two-photon calcium imaging in the brain slice preparation. Shown are neurons expressing jGCaMP8s in the cortical layer 2/3 superimposed a pair of stimulation electrodes. c, Stimulation protocol and representative neuronal responses evoked by AM_100_, AM_50_, and AM_0_ waveforms with 1Hz beat frequency. Each stimulus was applied for 1-s. Heat maps show responses of individual cells, and traces above indicate the average response across cells. d, Thresholds across AM conditions and carrier frequencies determined in individual neurons. Thresholds were quantified from 425 responsive neurons obtained from 12 slices across 5 mice. Violin plots show the distribution of thresholds; box plots indicate the median and interquartile range, and dots indicate individual neurons. Statistical significance was assessed using a LMM with FDR-corrected post hoc comparisons. \*\*\**p* < 0.001. e, Representative traces illustrating the onset of responses under different AM conditions. 2-kHz carrier frequency was used. The red hatched box indicates the timing of the stimuli. Three times the standard deviation of the baseline (3SD) was used to determine the latency used in the subsequent analysis. f, Latency of the onsets at 2-kHz. It was quantified in 140 cells from 5 mice and 12 slices. Violin plots show the latency distribution for each condition while box plots indicate the median and interquartile range and dots indicate individual neurons. Statistical significance was assessed using a LMM model with FDR-corrected post hoc comparisons. \*\*\**p* < 0.001.

We generated 1-s of two pmTI waveforms (AM_100_ and AM_50_) and one unmodulated high-frequency waveform (AM_0_) with varying current intensities and carrier frequencies. Individual neuron responses to pmTI stimulation were quantified as calcium transients observed in the brain slice preparation expressing genetically encoded calcium indicator, jGCaMP8s^43^, under two-photon scanning microscopy (Fig. 1b,c).

Across all carrier frequencies tested, all three AM indexes evoked neuronal activity (Fig. 1c). The amount of current required to elicit responses was consistently lowest with AM_0_ compared to stimulation with higher AM (Fig. 1d). A linear mixed-effects model (LMM) revealed significant effects of AM index (*F*(2,1266) = 3.38, *p* < 0.05) and carrier frequency (*F*(2,1266) = 29.71, *p* < 0.001), and the effect of AM index varied significantly across carrier frequencies (F(4,1266) = 2.91, p < 0.05). Thresholds were higher under AM50 and AM100 than under AM0 across carrier frequencies (all FDR-corrected *q* < 0.001). To test whether neuronal responses were driven by the abrupt onset of stimulation, we also applied stimulation with a gradual 30-s ramp-up to minimize rapid changes in stimulus amplitude. Neurons responded at similar stimulation amplitudes during both the no-ramp condition and the 30-s long-ramp-up condition (Supplementary Fig. 1).

Thresholds increased with carrier frequency, indicating that higher-frequency stimulation requires more current to evoke neuronal responses. Within the AM_0_ condition, thresholds increased significantly with the frequency of the carrier (FDR-corrected *q* < 0.001 in all comparisons). This frequency-dependent increase in threshold may reflect multiple factors, including frequency-dependent changes in tissue impedance and reduced neuronal responsiveness at higher carrier frequencies, as rapid field oscillations increasingly exceed the timescales of membrane and voltage-gated ion-channel dynamics.

We next analyzed the onset of responses using 2-kHz carrier frequency (Fig. 1e). While unmodulated high-frequency stimulation produced the fastest onset, pmTI stimulation delayed calcium transients, with the longest delay observed with AM_100_ (Fig. 1f), suggesting that action potential occurs when stimulation amplitude reaches a certain threshold. LMM revealed a significant effect of AM index on the latency (*F*(2,417) = 130.25, *p* < 0.001). Latencies for both AM_50_ and AM_100_ were significantly longer than for AM_0_ (FDR-corrected *q* < 0.001 in both comparisons), and latency for AM_100_ was significantly longer than for AM_50_ (FDR-corrected *q* < 0.001). Together, these results demonstrate that unmodulated high-frequency stimulation without AM alone is sufficient to evoke neuronal responses, with somewhat better efficacy than pmTI with AM.

### Amplitude modulated waveform enables rhythmic modulation of neuronal activity

The previous experiment used only a single cycle of stimulation and therefore did not address neuronal responses during multi-cycle TI stimulation. To address this, we applied 10-s pmTI stimulation consisting of 10 cycles of a 1Hz envelope.

Calcium transients revealed clear rhythmic fluctuations synchronized to the envelope frequency during AM_50_ and AM_100_ stimulation, whereas responses during AM_0_ lacked obvious periodicity (Fig. 2a). To quantify this, we calculated the power spectral density (PSD) from the calcium signals. The analysis revealed a clear peak at 1Hz for the AM_50_ and AM_100_ conditions, whereas no clear peak was seen with AM_0_. LMM model revealed a significant effect of AM index on the band-power between 0.8-1.2Hz (*F*(2,594) = 164.32, *p* < 0.001). Higher 1Hz power was found with higher AM (FDR-corrected *q* < 0.001 for AM_50_ *vs.* AM_0_, AM_100_ *vs.* AM_0_, and AM_100_ *vs.* AM_50_; Fig. 2b, c).

**Fig. 2.**
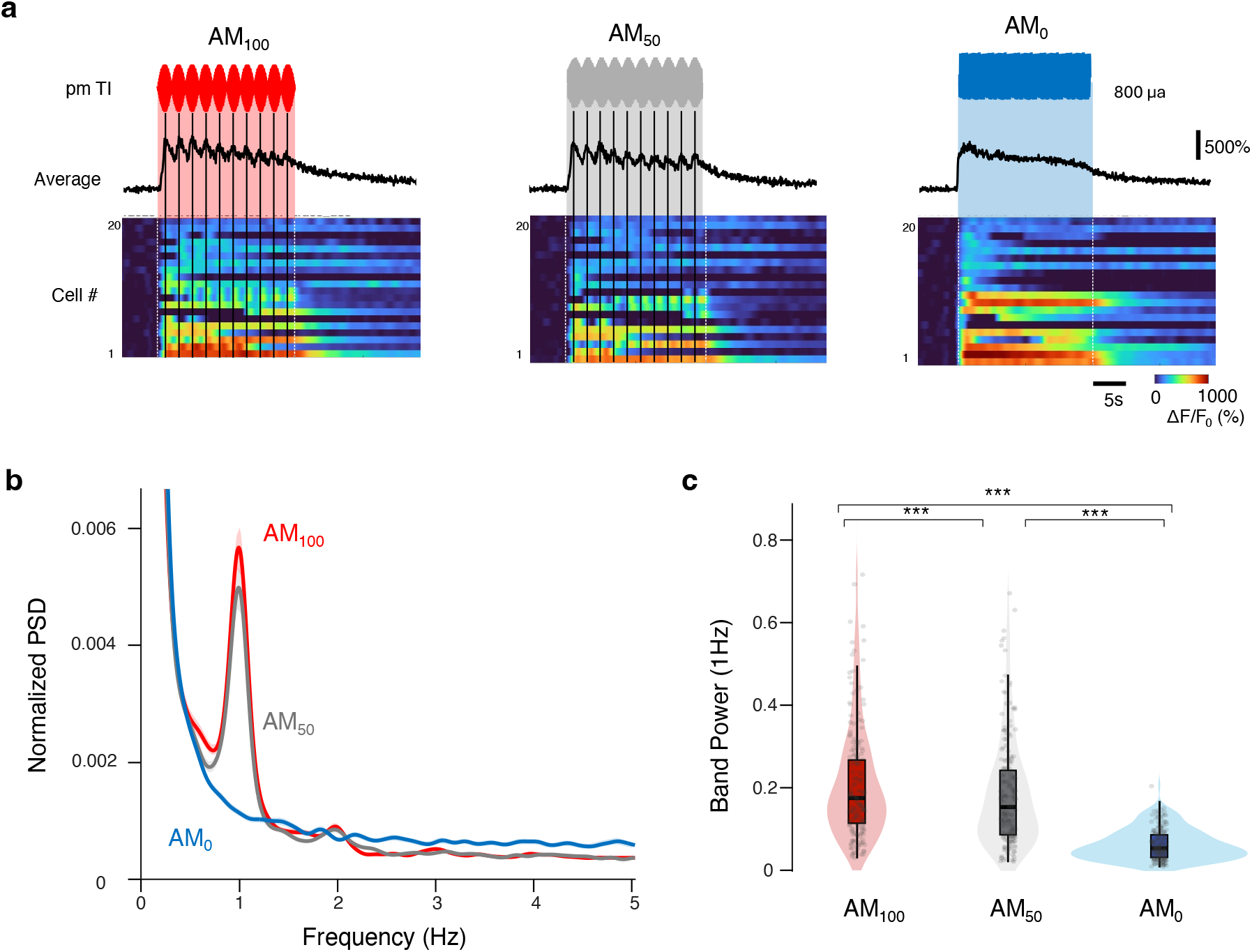
Rhythmic neuronal activity during prolonged pmTI stimulation. **a**, Representative calcium responses during 10-s pmTI stimulation at ±1mA. Heat maps show responses of individual cells, and traces above indicate the average response across cells. The peaks of AM envelopes are indicated by dotted line. b, PSD of calcium responses. Shaded lines indicate ±SEM. c, Quantification of the 1-Hz power. Band power between 0.8 and 1.2-Hz was used. Violin plots show the distribution of the bandpower for each condition while box plots indicate the median and interquartile range, and dots indicate individual neurons. Statistical significance was assessed using LMM model with FDR-corrected post hoc comparisons. \*\*\**p* < 0.001.

Although the slow kinetics of the calcium indicator does not allow precise quantification of spike counts within individual envelope cycles, these data showed that AM waveforms induce robust rhythmic activity aligned with the envelope frequency, which was absent in responses by unmodulated high-frequency stimulation. These findings indicate that AM is critical for imposing temporally structured neuronal activity.

### Computational modeling reveals ionic mechanisms underlying widespread activation and focal temporal modulation during TI stimulation

To investigate whether our experimental observations could be reproduced *in silico*, we first performed simulations using NEURON^44^ on 2 neuronal models: a previously published layer 2/3 cortical pyramidal neuron model^36,45,46^ and a reduced single-compartment model containing the ion channel composition of axons. These models incorporate various ionic conductance, including sodium and potassium channels with distinct kinetics (Fig. 3a).

**Fig. 3.**
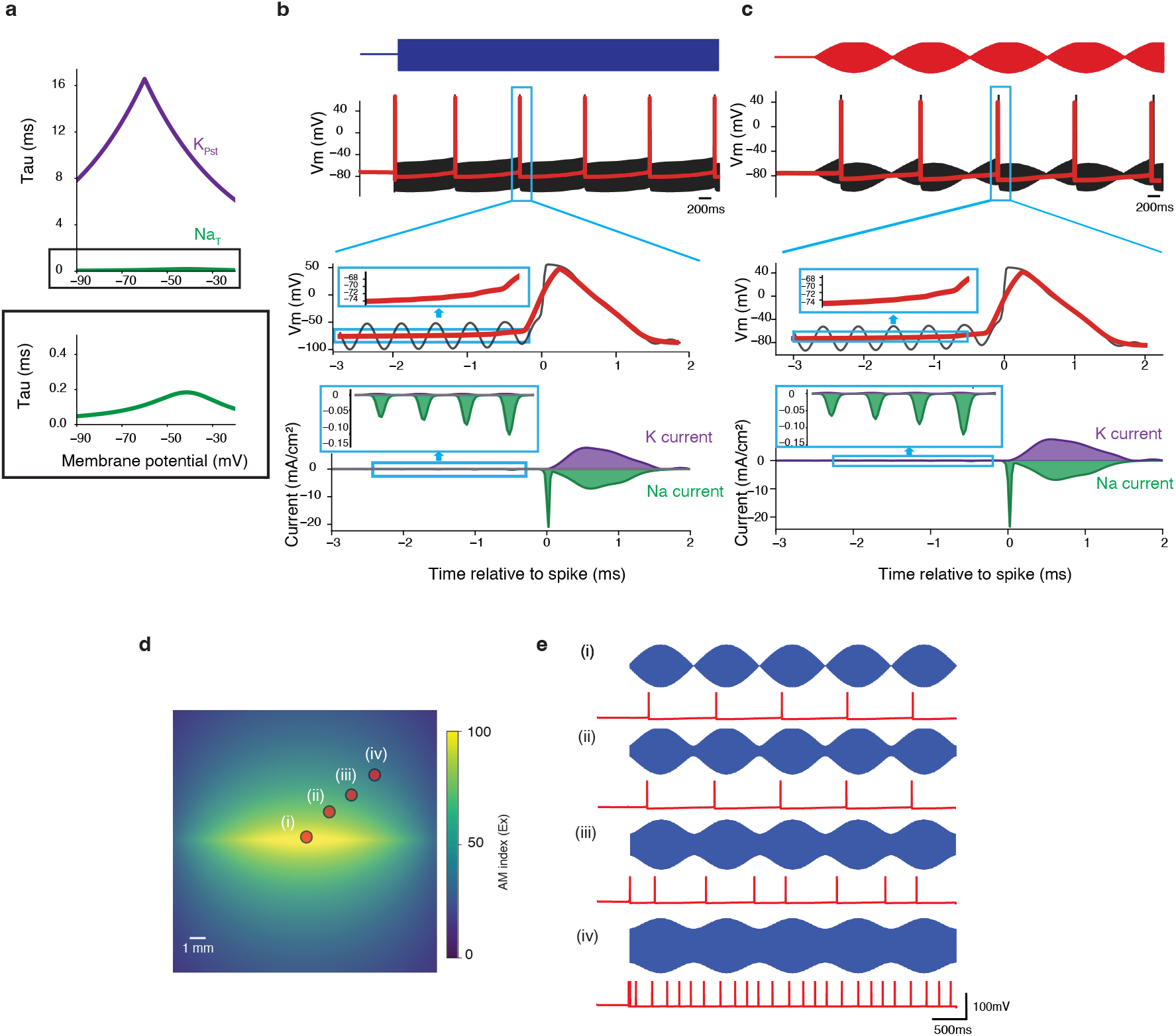
Computational modeling reveals ionic mechanisms underlying widespread activation and focal temporal modulation during TI stimulation. **a**, Simulated ionic mechanisms underlying spike generation during high-frequency stimulation. Voltage-dependent activation time constants (Tau) for sodium and potassium channels used in the neuronal model. Rapid sodium-channel activation and slower potassium-channel activation produced a transient inward current imbalance, resulting in gradual membrane depolarization before spike initiation. b, Simulated responses of cortical pyramidal neuron model to unmodulated high-frequency stimulation. Carrier-only stimulation elicited neural activation accompanied by progressive membrane depolarization before spike generation. Insets show membrane potential and ionic currents immediately preceding spike initiation. Purple represents potassium outflow. Green represents sodium inflow. c, Simulated responses of cortical pyramidal neuron model to amplitude-modulated stimulation. TI waveforms produced rhythmic responses synchronized to the modulation frequency. Insets show membrane potential and ionic currents immediately preceding spike initiation. d, Simulated conventinal TI stimulation. AM index map was generated by two electrode pairs delivering currents at 2000 and 2001 Hz. e, Representative simulated membrane potential traces from the four locations indicated in d (i– iv), corresponding to different AM index values within the TI field. Local electric field (blue) and membrane potential trace (red) are shown.

As we had performed in brain slices, we first applied premodulated waveforms to each model. Consistent with our experimental observations, both 2 models responded to both unmodulated (AM_0_) and 1Hz modulated (AM_100_) high-frequency stimulation (Fig. 3b-c, Supplementary Fig. 2). Although both AM_0_ and AM_100_ elicited neural activation, with multiple spikes occurring during stimulation, the spikes observed during AM_100_ appeared to synchronize to the 1Hz beat frequency. To investigate the underlying mechanism, we measured ionic currents immediately preceding spike initiation in each model. Across all models, membrane depolarization gradually accumulated because inward sodium current exceeded outward potassium current prior to spike generation. As a result, the membrane potential progressively increased at each cycle until it reached firing threshold. This phenomenon is likely attributable to differences in channel kinetics. Sodium channels activate rapidly in response to high-frequency stimulation, whereas potassium channels exhibit slower activation dynamics (Fig. 3a). Consequently, an imbalance emerges between inward sodium and outward potassium currents at each cycle, leading to a cumulative increase in the average membrane potential. Similar stimulation-induced depolarization has been reported in a previous experimental study, supporting the plausibility of this mechanism^27^.

Next, we examined how multiple neurons responded to conventional TI stimulation across space. To this end, a layer 2/3 cortical pyramidal neuron model was positioned throughout a two-dimensional field (2 cm × 2 cm), and TI stimulation was generated using two electrode pairs delivering currents at 2000 and 2001 Hz respectively, producing a 1Hz beat frequency (Fig. 3d). Importantly, the modified layer 2/3 cortical pyramidal neuron model showed that neuronal responses were observed throughout the field, with cells located near the electrodes often exhibiting higher firing rates due to a stronger local electric field (Fig. 3e). However, simulations revealed a clear spatial gradient in rhythmic modulation. Neurons located near the center of the TI field exhibited oscillatory responses at the beat frequency, whereas neurons closer to the stimulating electrodes showed less1-Hz modulation (Fig. 3e). These findings suggest that neuronal activation is broadly distributed across the field and is largely determined by local field strength. In contrast, synchronization to the beat frequency is spatially restricted and becomes strongest in regions with the highest AM index.

### TI stimulation produces widespread neuronal activation while preserving focal temporal modulation

We next tested the prediction from the model simulation above experimentally using TI stimulation. Two independent electrode pairs delivered sinusoidal currents at 2000 Hz only (carrier) or 2000 and 2001 Hz (TI), generating a 1Hz beat frequency within the cortex in brain slices (Fig. 4a, c). Each stimulation trial consisted of a 5-s stimulus with 0.5-s linear ramp-up and ramp-down periods to minimize the effect of stimulation onsets and offsets.

**Fig. 4.**
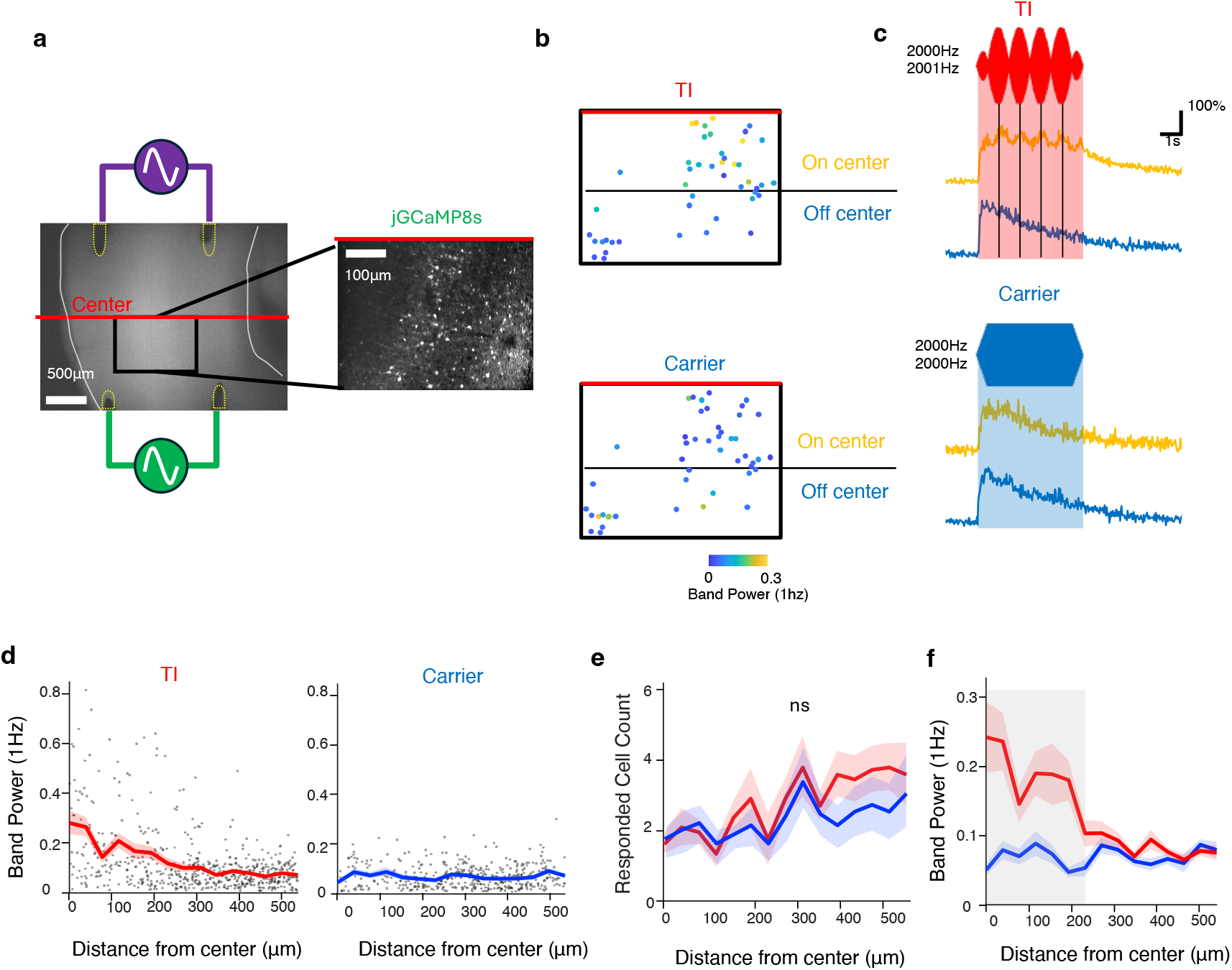
Temporally and spatially coordinated neuronal activity during TI stimulation. **a**, Experimental configuration. Two independent electrode pairs delivered sinusoidal currents at 2000 and 2001 Hz to generate a 1Hz beat frequency within the cortex of a brain slice. b, Spatial distribution of responsive neurons and beat-frequency modulation across the imaging field during TI and carrier stimulation at ±4mA. Power spectrum density was calculated for each cell and the band power between 0.8 and 1.2-Hz was measured and shown in pseudo color. c, Representative calcium traces during TI and carrier stimulation at ±4mA are shown for neurons located in half of the field of view closer to the interference centerline (On-center) and for those located in the other half farther from the centerline (Off-center). d, Spatial distribution of the 1Hz band power as a function of distance from the interference centerline during TI and carrier stimulation. Individual neurons (dots) were binned according to their distance from the interference field centerline to calculate averages (thick lines). The shaded lines indicate ±SEM. e, Responsive cell counts as a function of distance from the interference centerline. Shown are the mean value (thick lines) in each bin ±SEM (shaded lines). *n* = 667 and 555 responsive cells in 15 slices from 5 mice during TI and carrier stimulation, respectively. Statistical significance was assessed using a Poisson generalized LMM model. ns, not significant. f, Power at 1Hz as a function of distance from the interference centerline. Shown are the means (thick lines) of the band power between 0.8 and 1.2-Hz in the power spectrum density of calcium traces ±SEM (shaded regions). *n* = 667 and 555 responsive cells in 15 slices from 5 mice during TI and carrier stimulation, respectively. Statistical significance was assessed using a LMM model. A pairwise comparison at each bin (39.6 µm) showed significant increases in the 1Hz band power closer to the interference centerline (0, 39.6, 118.8, 158.4, and 198 µm from the centerline; FDR-corrected *p* <0.05, respectively).

We first examined the spatial distribution of neuronal activation across the imaging field. Consistent with our model, neurons responded to the stimulation throughout the field (Fig. 4b, d). To quantify this pattern, we counted responsive cells and binned them according to their distance from the interference field centerline. Poisson generalized LMM showed that the overall number of responsive cells did not differ significantly between TI and carrier stimulation (β = –0.003, p = 0.978). Moreover, TI did not significantly alter the spatial distribution of responsive cells across distance from the interference field centerline (β = 0.345, p = 0.057), indicating that neural recruitment itself was not spatially restricted to the predicted TI target region (Fig. 4d, e, Supplementary Fig. 3).

We next investigated whether temporal modulation exhibited a distinct spatial profile. Neurons located close to the interference centerline displayed clear rhythmic activity synchronized to the 1 Hz envelope, whereas neurons located farther from the center showed weaker or absent oscillatory modulation despite remaining responsive to stimulation (Fig. 4b-d). To quantify this effect, we analyzed the band power around 1 Hz from the power spectrum density of calcium traces. The 1 Hz synchronization power was significantly higher during TI than during carrier stimulation (F(1,1218) = 175.71, p < 0.001) (Fig. 4d, f). This TI-associated increase of 1 Hz power became progressively weaker as distance from the field center increased (F(1,1218) = 106.18, p < 0.001; β = –0.176). This spatial pattern closely matched the predicted distribution of AM index (Supplementary Fig. 3) and was consistent with the computational modeling results. These data reveal a dissociation between overall neuronal activity levels and temporal modulation during TI stimulation. TI stimulation does not confine neuronal activation to the target. Rather, the TI stimulation appears to selectively enhance temporal modulation near the interference center.

### TI stimulation induces rhythmic neural activity in anesthetized mice

To determine whether the temporal modulation observed in slice preparations generalizes to intact brain, we performed *in vivo* two-photon calcium imaging while applying TI stimulation targeting the left motor cortex (Fig. 5a). TI stimulation was delivered for 10-s with 0.5-s linear ramp-up and ramp-down periods under anesthesia to minimize spontaneous activity. The stimulation configuration was designed to maximize AM at the center of the cranial window. The presence of AM was also confirmed even at the edge of the cranial window, away from the center, using an EEG electrode placed near the window (Fig. 5b).

**Fig. 5.**
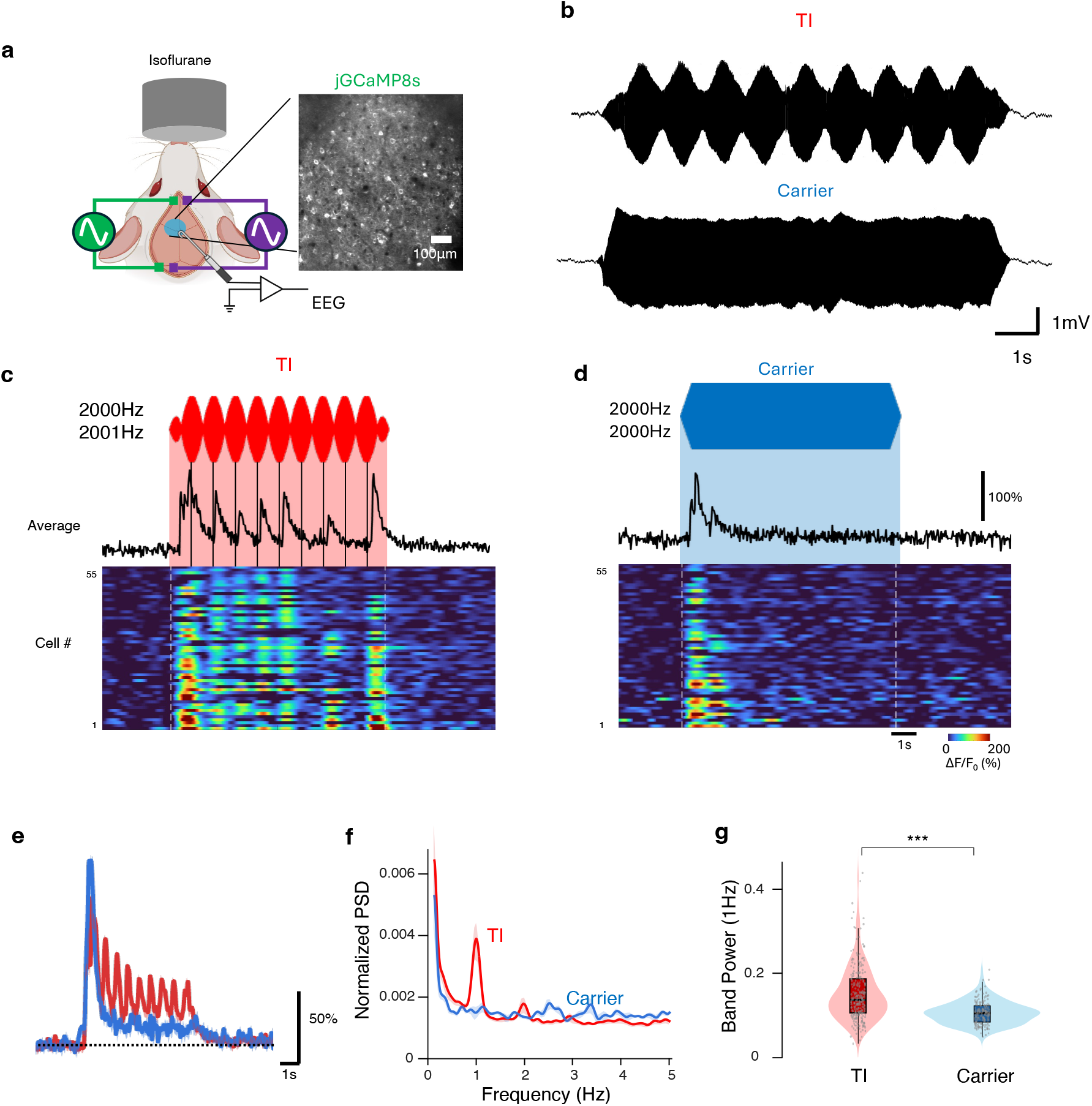
TI stimulation induces rhythmic activity in anesthetized mice. **a**, Experimental configuration for *in vivo* two-photon calcium imaging. TI stimulation was delivered through two pairs of screws chronically implanted in the skull under isoflurane anesthesia. A chronic cranial window was placed over the cortex between the skull screws. A epidural electrode was placed at the edge of the window to record stimulation waveforms. b, Representative waveforms recorded during TI stimulation and carrier stimulation at ±1mA using an EEG electrode positioned at the edge of the cranial window. c, Representative calcium transients during TI stimulation at ±1mA (red shaded rectangle). Heat maps show responses of individual cells, and traces above indicate the average response across cells. Dotted lines were added to highlight the envelope peaks. d, Representative calcium transients during carrier stimulation alone at ±1mA (blue shaded rectangle) in the same mouse. Heat maps show responses of individual cells, and traces above indicate the average response across cells. e, Calcium transients across all the responsive cells. Shown are means (thick traces) ± SEM (thin traces, *n* = 252 cells for TI (red) and *n* = 178 cells for carrier stimulation alone (blue) from 5 mice. f, Normalized PSD of calcium transients. PSD was calculated for individual cell traces and then averaged (thick traces). The same data set as in the previous panel was used. Thin traces represent SEM. g, Quantification of 1-Hz band power. Violin plots show the distribution for each condition; box plots indicate the median and interquartile range, and dots indicate results from individual cells. LMM, n = 252 cells for TI and *n* = 178 cells for carrier stimulation alone from 5 mice, \*\*\**p* < 0.001.

TI stimulation elicited robust neuronal responses observed as rhythmic calcium transients synchronized to the 1Hz beat frequency (Fig. 5c), indicating that frequency-specific modulation was preserved *in vivo*. High-frequency stimulation alone also induced neuronal activity of comparable amplitude (Fig. 5d, e). However, this activity lacked a corresponding low-frequency oscillatory component. Power spectral analysis further supported this distinction. Under TI stimulation, neuronal activity exhibited a distinct peak at 1Hz, whereas this peak was absent under 2000Hz carrier stimulation alone (Fig. 5f, g). LMM revealed 1Hz band power was significantly higher during TI stimulation than during carrier only stimulation (LMM, *F*(1,428) = 76.07, *p* < 0.001). These results are similar to those obtained in slice preparations, demonstrating that *in vivo* TI stimulation can impose beat frequency-specific modulation to neuronal activity and that high-frequency stimulation alone can also induce activity of similar magnitude, albeit with a lack of temporal modulation. To determine whether this rhythmic modulation was specific to a 1Hz beat frequency, we additionally tested TI stimulation with a 0.5-Hz beat frequency. TI stimulation at 0.5-Hz produced neuronal responses with a corresponding 0.5-Hz rhythmic component, with increased power at the applied beat frequency (Supplementary Fig. 4). These findings indicate that TI-induced rhythmic modulation is not restricted to 1Hz, but can follow the applied beat frequency.

### Prolonged TI stimulation induces expression of an immediate-early gene at the off-target region

While two-photon imaging provides high-resolution measurements of neuronal activity in individual cells, it cannot assess activity outside the field of view. We therefore conducted an immunohistological analysis using c-Fos, an immediate-early gene whose expression levels reflect recent neuronal activity^10,47–49^ to examine neuronal activation across cortical and deep brain regions following prolonged TI stimulation. As a comparison, we also applied high-frequency stimulation but of the same frequency (carrier only). We first confirmed that neuronal responses were sustained during prolonged stimulation using two-photon calcium imaging in the same mice used for the experiments in Fig. 5. For prolonged stimulation, the 10-s on, 10-s off stimulation paradigm was applied for 20 minutes. Both TI and carrier-only stimulation elicited responses throughout the stimulation period, although response amplitudes gradually decreased over time (Supplementary Fig. 5).

A similar prolonged stimulation paradigm was used to examine the spatial extent of neuronal activation. Stimulation electrodes were arranged on the skull so that AM occurred within the hippocampus, as described previously^50,51^. We were able to record strong AM in the hippocampus in several mice (Fig. 6a). In a separate cohort of animals, we then quantified c-Fos expression in the cortex, and in the CA1 layer and dentate gyrus (DG) granule cell layer of the hippocampus following 20 minutes of stimulation. All mice underwent anesthesia and electrode placement and received either TI stimulation, carrier only stimulation or no stimulation (sham) control). Compared with sham control, the number of neurons expressing c-Fos was significantly increased by TI stimulation in the non-targeted cortex below the stimulation electrodes, whereas no such increase was detected within the target area, including CA1 and DG (Fig. 6b, c). Similarly, significant c-Fos upregulation was observed in the cortex, but not in the hippocampus under the carrier-only stimulation condition (Fig. 6b, c).

**Fig. 6.**
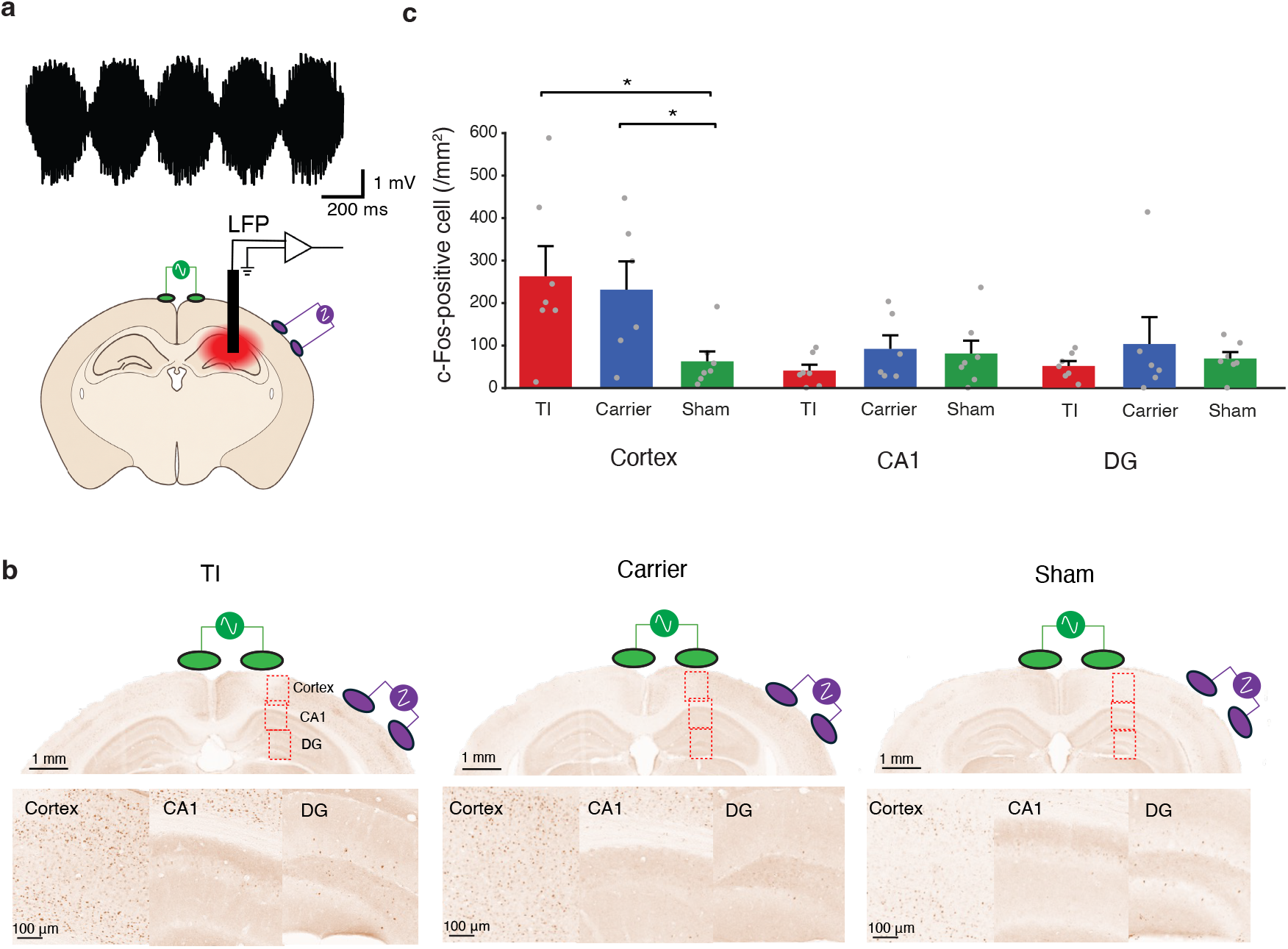
c-Fos immunolabeling reveals off-target cortical activation during prolonged TI stimulation. **a**, Representative recording of AM electric field in the hippocampus during TI stimulation with ±400 μA at 2000 and 2005 Hz, together with a schematic of the configurations of the stimulation and the recording electrodes. Green and purple ovals indicate the two independent electrode pairs used to deliver the kilohertz currents. A black rectangle denotes a recording electrode for local field potential (LFP). b, Representative coronal sections showing c-Fos immunohistochemistry in the cortex, CA1, and dentate gyrus (DG) following 20 minutes of TI stimulation with ±500 μA at 2000 and 2010 Hz (10-s on/10-s off; left), carrier-only stimulation with ±500 μA at 2000 Hz (10-s on/10-s off; middle) or no stimulation (sham control; right). The ovals indicate the approximate locations of the two stimulation electrode pairs. c, Quantification of c-Fos-positive cells in the cortex, CA1, and DG. Bars show mean + SEM. *n* = 7 mice (TI), *n* = 6 mice (carrier-only), and *n* = 7 mice (sham). Statistical significance was assessed using one-way ANOVA followed by pairwise t-tests with Holm–Bonferroni correction. \**p* < 0.05.

These findings show that the prolonged kilohertz-frequency stimulation can induce significant neuronal activation in the cortex, where the local electric-field magnitude is high, regardless of the presence of AM. Despite the clear indication of AM in the target, c-Fos expression did not significantly increase in the hippocampus under the prolonged TI stimulation condition, suggesting that the local electric-field magnitude was insufficient to drive the expression of this immediate-early gene. These results indicate that neuronal activation, as reflected by c-Fos expression, is more closely associated with the local electric-field magnitude of kilohertz-frequency stimulation than with the presence of AM.

### Amplitude modulated waveform stimulation induces rhythmic subthreshold depolarization

The experiments thus far were based on calcium imaging or histological analyses, a technique that primarily relies on suprathreshold spiking in neurons^43^ and, thus, do not necessarily mimic the effect of TI when the stimulus amplitude is subthreshold. We therefore investigated how subthreshold TI stimulation modulates the membrane potential in slice preparations expressing a genetically encoded voltage indicator (GEVI), JEDI-2P (Fig. 7a), which can report both suprathreshold and subthreshold changes in the membrane potential^52^. We used pmTI stimulation (AM_100_) and carrier-only stimulation (AM_0_) with either ±250 or ±500 µA, which we deemed subthreshold for the majority of the cells based on the first set of experiments (Fig. 7a).

**Fig. 7.**
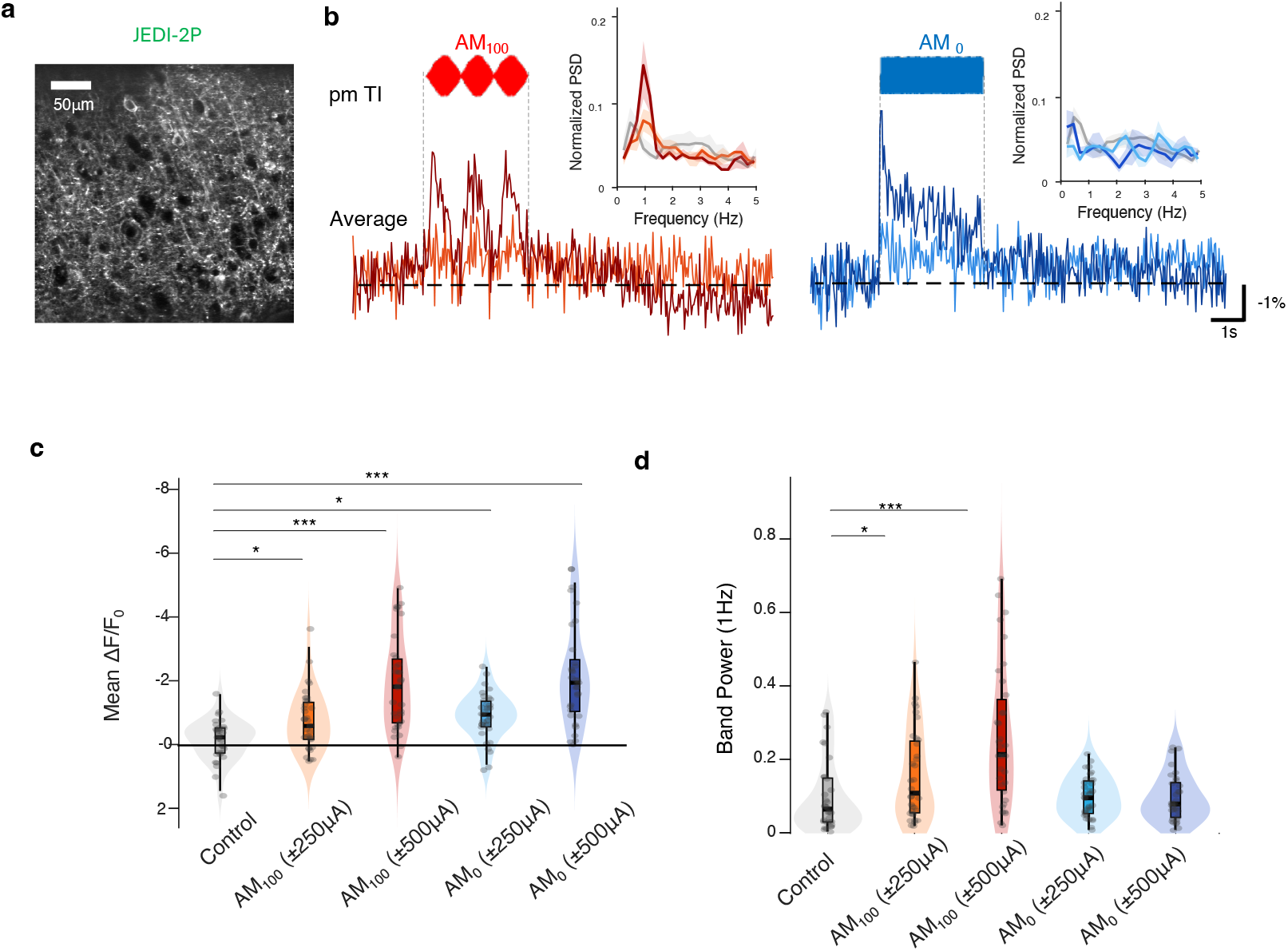
pmTI induces rhythmic subthreshold depolarization. **a**, Representative fluorescence image from an acute brain slice expressing JEDI-2P. b, Representative GEVI transients during subthreshold stimulation with pmTI stimulation (AM_100_) and carrier-only stimulation (AM_0_). Traces show the global average fluorescence signal in the imaging field during stimulation, and insets show the corresponding normalized PSD. An upward change represents a decrease in fluorescence and an increase in membrane potential. c, Means of relative change in fluorescence during stimulation. Violin plots show the distribution for each condition; box plots indicate the median and interquartile range, and dots indicate individual trials. Statistical significance was assessed against the control where no stimulation was applied using LMM, followed by FDR correction. *n* = 33 trials for control, 35 for AM_100_ (±250 µA), 35 for AM_100_ (±500 µA), 35 for AM_0_ (±250 µA), and 33 for AM_0_ (±500 uA), from 8 slices across 4 mice. \**p* < 0.05, \*\*\**p* < 0.001. d, Normalized power at 1 Hz during stimulation. Violin plots show the distribution for each condition; box plots indicate the median and interquartile range, and dots indicate individual trials. Statistical significance was assessed using LMM, followed by FDR correction. Normalized 1 Hz power was significantly increased relative to baseline for AM_100_ (±250 uA) and AM_100_ (±500 uA), whereas neither AM_0_ (±250 uA) nor AM_0_ (±500 uA) showed a significant change. n = 33 trials for control, 35 for AM_100_ (±250 uA), 35 for AM_100_ (±500 uA), 35 for AM_0_ (±250 uA), and 33 for AM_0_ (±500 uA), from 8 slices across 4 mice. \**p* < 0.05, \*\*\**p* < 0.001.

GEVI recordings showed that both pmTI stimulation (AM_100_) and carrier-only stimulation (AM_0_) produced depolarization (a decrease in fluorescence) during stimulation (Fig. 7b, c). The mean fluorescence during stimulation was significantly different from baseline (a period before stimulation) in all four stimulation conditions (LMM, *F*(4,166) = 27.39, *p* < 0.001; FDR-corrected q < 0.05, < 0.001, < 0.05, and < 0.001 for AM_100_ with ±250 µA, AM_100_ with ±500 µA, AM_0_ with ±250 µA, AM_0_ with ±500 µA, respectively; Figs. 7b, c), indicating that both pmTI and carrier-only stimulation induced subthreshold depolarization that were not seen using calcium imaging. We then examined whether these responses contained rhythmic modulation at the beat frequency. Under pmTI stimulation (AM_100_), the GEVI signal exhibited clear oscillatory dynamics synchronized to the 1 Hz envelope, and this rhythmic component became larger at higher stimulation intensity (Fig. 7c, d). In contrast, carrier-only stimulation (AM_0_) did not produce such oscillatory modulation. Normalized power at 1Hz during stimulation was only significant in the pmTI conditions (LMM, *F*(4,166) = 9.90, *p* < 0.001; FDR-corrected *q* < 0.05 and < 0.001 for 250µA and 500µA, respectively) but not in the carrier-only conditions (FDR-corrected *q* = 0.798 for both). These results indicate that high-frequency stimulation can induce depolarization whereas pmTI stimulation can elicit rhythmic modulation at the beat frequency even when neurons are not firing action potentials.

### TI stimulation induces rhythmic activity in awake mice

We next examined the impact of TI stimulation in awake, head-fixed, but otherwise freely moving mice (Fig. 8a). We confirmed that brief TI stimulation at ±1 mA with the 1-Hz beat frequency targeting the left motor cortex produced visible movement of the contralateral forelimb, consistent with previously reported motor responses to TI stimulation^10^. Unlike anesthetized conditions where spontaneous spiking was almost absent, we observed occasional spontaneous activity through the imaging window. Each stimulation consisted of a 30-s stimulation period with 0.5-s ramp-up and 0.5-s ramp-down phases. We did not induce direct responses when stimulation amplitude was ±250 µA under anesthesia (Supplementary Fig. 6). Therefore, two amplitudes (±50 and ±200 μA) were tested as subthreshold level stimulations.

**Fig. 8.**
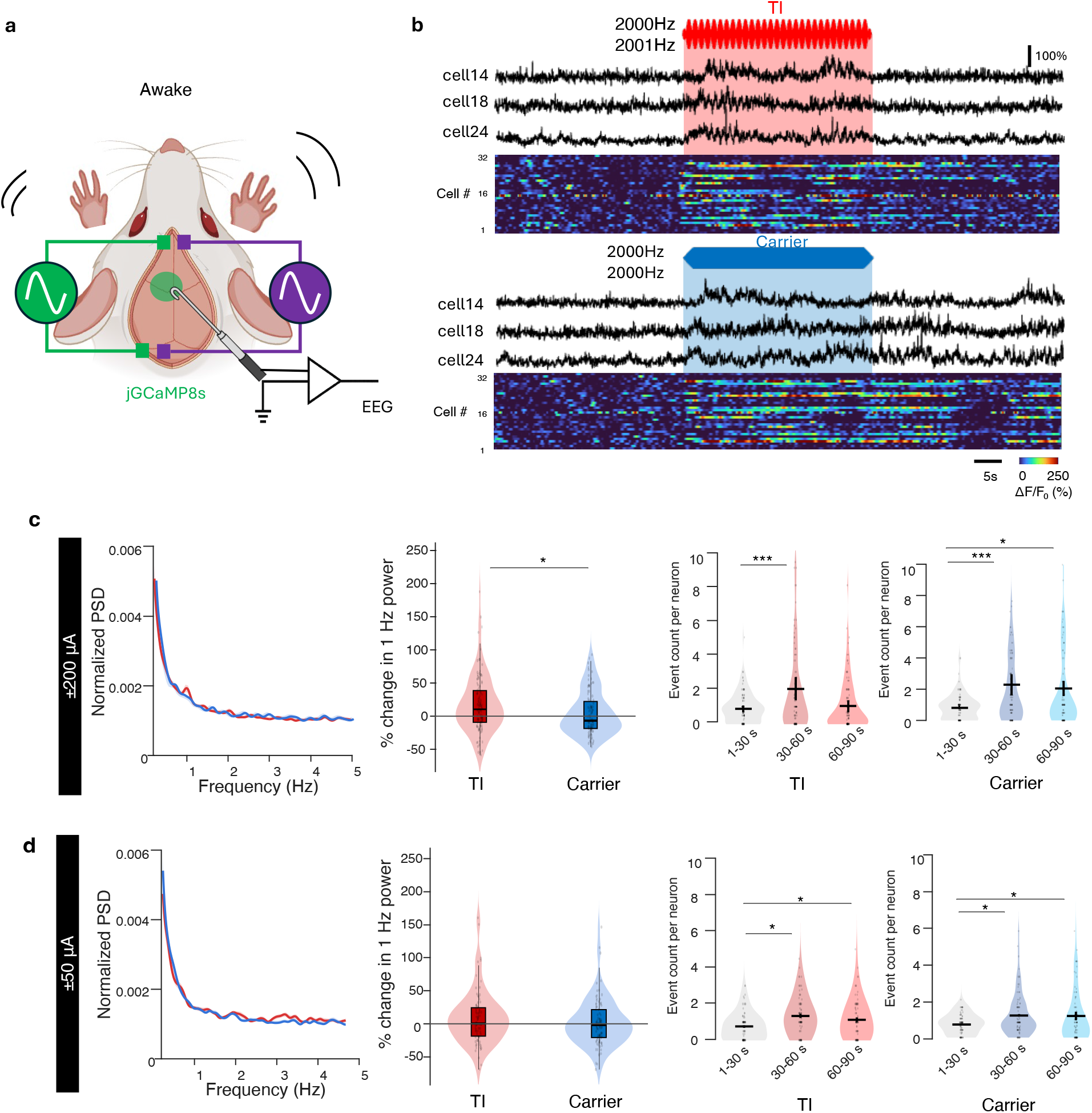
TI-induced rhythmic activity in awake mice. **a**, Experimental configuration for *in vivo* calcium imaging during TI stimulation in awake, head-fixed, but otherwise freely moving mice. b, Representative calcium change during TI and carrier stimulation at ±200 μA and ± 50μA. Traces show representative calcium transient from responsive neurons at ±200 μA and heat maps show responses from 32 neurons in the field. The shaded rectangles indicate the stimulation periods. c, Neuronal activity at ±200 μA. Left, normalized PSD during TI and carrier stimulation. Middle, quantification of 1-Hz power. Right, event counts before, during, and after stimulation for TI and carrier conditions. Each dot represents one neuron with repeated observations from the same neuron averaged within each condition. Violin plots show the distribution for each condition; box plots indicate the median and interquartile range, and dots indicate individual neurons. *n* = 107 and 102 neurons from 3 mice for TI and carrier only condition, respectively. Statistical significance was assessed using LMM. \**p* < 0.05. d, Neuronal activity at ±50 μA. Left, normalized PSD during TI and carrier stimulation. Middle, quantification of 1 Hz power. Right, event counts before, during, and after stimulation for TI and carrier conditions. Each dot represents one neuron, with repeated observations from the same neuron averaged within each condition. Violin plots show the distribution for each condition; box plots indicate the median and interquartile range, and dots indicate individual neurons. *n* = 82 and 89 neurons from 3 mice for TI and carrier only condition, respectively. Statistical significance was assessed using LMM. \**p* < 0.05. \*\*\**p* < 0.001

At ±200 μA, we observed clear oscillatory modulation at beat frequency (1-Hz) during TI stimulation (Fig. 8b), which produced a significantly greater increase in 1-Hz power than carrier alone as revealed by LMM (*F*(1,207) = 5.00, *p* < 0.05; Fig. 8c). In contrast, at ±50 μA, LMM showed that TI stimulation did not produce a significant increase in 1-Hz power compared to carrier alone (*F*(1,169) = 0.85, *p* = 0.357). We further analyzed the number of calcium transients during stimulation. During TI stimulation, event counts increased significantly at both ±50 μA (*F*(2,243) = 8.37, *p* < 0.001) and ±200 μA (*F*(2,318) = 13.33, *p* < 0.001). Likewise, high-frequency stimulation alone also increased neuronal activity during stimulation at both amplitudes (*F*(2,261) = 8.40, *p* < 0.001 and *F*(2,288) = 17.26, *p* < 0.001 for ±50 µA and ±200 µA, respectively), but it did not produce significant beat-frequency oscillatory modulation. Thus, low-amplitude TI stimulation increased neuronal activity without clear entrainment, whereas high-amplitude TI stimulation induced both neuronal activation in the imaging field and rhythmic modulation at beat frequency in awake mice.

We additionally tested TI and carrier stimulation using a 5000Hz carrier at ±200 μA (Supplementary Fig. 7). Similar to the 2000Hz conditions, both TI and carrier stimulation paradigms increased neuronal activity during stimulation. However, TI stimulation at 5000Hz did not significantly increase 1Hz power at this amplitude. Together with the carrier-frequency dependence observed in our slice experiments, these results suggest that higher carrier frequencies may require greater stimulation amplitudes to produce measurable entrainment.

## Discussion

In this study, we systematically characterized neural responses to temporal interference (TI) stimulation across computational modeling, *ex vivo* preparations, and *in vivo* imaging at individual cellular levels. We found that unmodulated high-frequency stimulation alone was sufficient to evoke neuronal activity and that unmodulated kilohertz-frequency stimulation is not physiologically inert (Fig. 1-8). Although both unmodulated high-frequency stimulation and AM waveforms induced neural responses, AM played a distinct role in shaping the temporal structure of neuronal activity (Fig. 2-5,7,8, Supplementary Fig. 4). Across *ex vivo* and *in vivo* experiments, as well as computational models, TI stimulation consistently generated neural activity entrained to the beat frequency, whereas carrier-only stimulation produced neuronal activity without a distinct temporal pattern. Importantly, neuronal activation was broadly distributed across the field of view, whereas beat-frequency entrainment remained spatially localized to regions of AM modulation, the center region surrounded by the two pairs of electrodes (Fig. 4). These findings suggest that the primary effect of TI is not focal neural activation, but focal temporal modulation. TI-induced rhythmic activity was observed even as subthreshold membrane depolarization, indicating that temporal modulation can occur even when stimulation is not evoking action potentials (Fig. 7,8).

A central finding of this study is that neurons are sensitive to kilohertz-frequency electric fields. Both our experimental observations and computational modeling demonstrated that unmodulated high-frequency stimulation alone was sufficient to evoke neural responses, indicating that neuronal activation during TI cannot be explained solely by extraction of the low-frequency envelope. Instead, our simulations support a potential mechanism in which differences in the kinetics of sodium and potassium channel produce a progressive accumulation of membrane depolarization during high-frequency stimulation, consistent with previous studies implicating sodium-channel dynamics in high-frequency stimulation-induced depolarization^53^. Because sodium channels are activated more rapidly than potassium channels, each stimulation cycle generates a transient excess of inward current that is only partially opposed by delayed potassium currents. Even if neurons cannot generate action potential at individual kilohertz cycles, these small imbalances accumulate over time, progressively depolarizing the membrane until spike threshold is reached. Neural responses during both carrier-only and TI stimulation can be explained by this same mechanism^34^. In both conditions, neurons are driven primarily by the same high-frequency electric field and therefore undergo the same progressive depolarization. The key distinction is that, during TI stimulation, the amplitude of the high-frequency field varies according to the beat envelope. Consequently, membrane depolarization also fluctuates over time, producing rhythmic changes in neuronal excitability that give rise to beat-frequency entrainment. These findings suggest that the principal role of amplitude modulation is not to enable neuronal activation, but to organize the timing of activity generated by the underlying high-frequency stimulation.

Across our experiments, this entrainment increased with the strength of AM, suggesting that the AM index is a physiologically meaningful predictor of modulation strength. Importantly, we further demonstrated that TI can induce rhythmic membrane depolarization even under subthreshold conditions, indicating that beat-frequency modulation can emerge without directly evoking action potentials. However, the presence of subthreshold modulation does not necessarily imply that arbitrarily weak stimulation is sufficient to entrain neural activity *in vivo*. Previous studies have suggested that TI may alter spike timing without producing robust rhythmic firing^14^, and our findings support this and suggest a potential explanation for these observations. In our recordings in awake mice, lower stimulation amplitudes produced neural responses but failed to generate clear beat-frequency entrainment, whereas stimulation with higher amplitudes induced robust rhythmic neuronal activity. Similar observations were reported in conventional tACS studies where entrainment to externally imposed oscillations increase with stimulation intensity^54,55^. The entrainment appears as competition between external stimuli and ongoing endogenous network dynamics^56^. Thus, for TI to produce measurable entrainment, TI-induced modulation needs to overcome intrinsic fluctuations. Notably, as demonstrated in our experiments, higher-frequency carriers required substantially larger stimulation amplitudes to produce comparable physiological effects, raising the possibility that effective TI entrainment may require stronger stimulation than is commonly assumed. Kilohertz-frequency stimulation is generally well tolerated in humans^39^, suggesting that effective TI entrainment may still be achievable within practical stimulation ranges.

At the same time, effective TI modulation is unlikely to depend solely on stimulation intensity. Neural circuits possess intrinsic oscillatory dynamics that may be amplified by externally imposed rhythms. The efficacy of non-invasive brain stimulation is known to depend on behavioral state and ongoing sensory or cognitive processes^57–60^. Consequently, TI-induced entrainment may be enhanced when stimulation is paired with specific behavioral states, sensory inputs, or endogenous neural oscillations. Future studies should therefore investigate the range of stimulation amplitudes that can safely and reliably generate meaningful entrainment. They should also leverage the circuit and behavioral conditions under which TI-induced modulation can be most effectively amplified.

Interestingly, previous human fMRI studies reported robust BOLD responses in target regions receiving strong temporal interference modulation but did not detect significant cortical BOLD changes under the stimulation electrodes^15–18,24^. Our experiment demonstrated that kilohertz stimulation induced membrane depolarization and increased neuronal activity even in the absence of AM. Consistent with our findings, previous human studies have also reported that low-kilohertz transcranial alternating current stimulation increases cortical excitability^32^, suggesting that kilohertz electric fields can alter neuronal function despite the absence of detectable BOLD responses. One possible explanation for this discrepancy is the distinct relationship between neuronal activity and the BOLD signal. Previous neurophysiological studies have suggested that the BOLD signal is more closely related to local synaptic population activity, reflected by local field potentials, than to spike rate alone^61–63^. Therefore, off-target neuronal activation may not produce detectable BOLD responses despite increased neuronal excitability if the activity lacks sufficient temporal coordination. In addition, neuronal responses to sustained unmodulated kilohertz stimulation may undergo adaptation at the subthreshold level over time. In contrast, the periodic amplitude modulation of TI stimulation may maintain or repeatedly recruit coordinated neural responses at the beat frequency, thereby producing a more robust hemodynamic response in target regions.

Taken together, our findings suggest that TI stimulation may be more appropriately viewed as a combination of widespread neural activation and localized temporal modulation. Under this framework, high-frequency electric fields broadly influence neuronal excitability throughout the stimulated region, whereas amplitude modulation selectively imposes rhythmic organization on neural activity within regions of high AM index. This framework has several important implications for future studies of TI stimulation. First, if widespread activation is detrimental, stimulation strategies that preserve strong amplitude modulation within the target region while reducing activation elsewhere can be employed. For example, multi-pair stimulation paradigms may improve the balance between localized temporal modulation and off-target neural activation^25,64,65^. Second, widespread activation during TI stimulation can be exploited therapeutically. Previous studies using TMS and other neuromodulation approaches have demonstrated that modulation of connected cortical regions can indirectly influence distant targets through network interactions^66–68^. Similarly, TI stimulation may provide an opportunity to combine activation of selected cortical circuits with localized rhythmic modulation of deeper brain structures. Rather than considering only where amplitude modulation is generated, future studies should evaluate how both widespread activation and localized temporal modulation contribute to therapeutic outcomes across different neurological and psychiatric disorders.

Beyond its implications for TI, our findings suggest that kilohertz-frequency stimulation itself may provide a useful regime for neuromodulation, particularly given its relatively mild skin sensation^39^. Our experiments demonstrate neuronal activation through unmodulated kilohertz stimulation. Although we mostly focused on responses during stimulation, previous studies have reported effects that persist beyond the stimulation period^32,37^, suggesting that kilohertz stimulation could potentially provide a means of enhancing neural excitability for therapeutic neuromodulation. Furthermore, our pmTI experiments demonstrate that neuronal responses can be temporally modulated by AM waveforms. In superficial regions such as the cortex, this approach could potentially be delivered using a single electrode pair rather than two interfering pairs to introduce stimulation dynamics aligned with endogenous neural rhythms, such as alpha or beta activity.

Several limitations should be considered when interpreting these findings. First, our experiments were performed primarily in *ex vivo* brain slices and *in vivo* mouse cortex, and it remains unclear how these mechanisms generalize across cell types, brain regions, and brain sizes. Second, the kinetics of calcium indicators limit the temporal resolution with which TI-induced entrainment can be resolved at the level of individual cells; higher temporal resolution will be needed to further characterize spike-level entrainment. Moreover, while the entrainment specific to envelope frequency was observed at both 0.5 and 1 Hz (Figs. 2-7, Supplementary Fig. 4), systematic characterization across a broader range of envelope frequencies will be important for defining the frequency dependence. Third, kilohertz electrical currents might stimulate the peripheral nerves and/or muscles that could have contributed to the observed neural activity in the brain, especially in awake mice. However, neuronal activity and limb movement locked to the 1-Hz beat frequency suggests that these secondary effects are unlikely to fully account for the responses observed in the brain. Future studies with simultaneous behavioral monitoring will help further distinguish the primary and the secondary effects of TI stimulation. Finally, our results indicate that high stimulation amplitudes are necessary to achieve robust entrainment. Whether such modulation can be safely and effectively achieved in humans remains an important question for future translational studies.

Our results suggest that TI stimulation should be viewed not as a method for producing focal neural activation, but as a method for producing focal temporal modulation with widespread activation. We demonstrated that high-frequency stimulation itself broadly influences neuronal activity, whereas amplitude modulation selectively shapes the timing of that activity. This work provides a new perspective on how TI interacts with neural circuits and how its effects should be interpreted. Moving forward, exploiting localized temporal modulation jointly with widespread activation may enable more effective applications of TI, while the ability of kilohertz-frequency stimulation itself to drive neural activity may provide new opportunities for brain stimulation.

## Method

### Animals

All protocols involving animals were in accordance with Emory Institutional Animal Care and Use Committee. B6 albino mice (Charles River strain code: 493) were bred in house. They were kept in 12-hour dark/12-hour light cycles in a climate-controlled room with *ad libitum* access to regular rodent chow and water.

### Viral injections

Intracerebroventricular (ICV) injections of recombinant adeno-associated viral (rAAV) vectors were performed in neonatal pups to express a calcium indicator (rAAV9/hSyn-soma-jGCaMP8s; Addgene #: 169258; a gift from Marianne Fyhn) or a voltage indicator (rAAV9/hSyn-JEDI-2P-Kv-WPRE; Addgene #: 179463; a gift from Francois St-Pierre). Postnatal day 0 or 1 pups were removed from the dam, and anesthesia was induced by hypothermia by placing pups on a metal surface in an ice bucket until the movement stopped completely. We placed a sheet of Kim wipes between the pups and the metal to prevent frostbite. The dorsal surface of the head was visualized under a light microscope, and the skin was cleaned using alcohol pads. A microsyringe (10 µL, Gastight Neuros Syringe, Small RN, PN: 65460-06, Hamilton Company, Franklyn, MA) with the stopper set at 2 mm was manually lowered into the lateral ventricle as described^69^. A volume of 1 µL of viral particles (∼1×10^10^ viral genomes by qPCR for either AAV) diluted in phosphate-buffered saline (PBS) containing 0.05% trypan blue was slowly injected into each hemisphere. Following injection, pups were placed onto a warming pad until normal skin color and spontaneous movement were restored and returned to the dam. We left at least one pup in a litter to the dam all the time to minimize maternal rejection.

### Brain slice preparation

*Ex vivo* experiments were performed using mice that had received neonatal ICV injections. Animals of both sexes were used without distinction and were sacrificed for slice electrophysiology experiments between postnatal day 18 and 30. Prior to decapitation, mice were deeply anesthetized with 5% (v/v) isoflurane in oxygen followed by ketamine administration (200 mg/kg). Thereafter, the brain was immediately removed by dissection placed in ice-cold cutting solution (in mM) 87 NaCl, 25 NaHCO_3_, 2.5 KCl, 1.25 NaH_2_PO_4_, 2 MgCl_2_, 1 CaCl_2_, 10 glucose, and 75 sucrose. Brain slices (250 μm) were sectioned in the coronal plane using a vibrating blade microtome (VT1200S, Leica Biosystems, Richmond, IL) in the same solution. Slices were transferred to an incubation chamber and maintained at 34°C for ∼60 min and then at room temperature (23–24°C) thereafter.

During recordings, a slice was placed into an imaging chamber continuously superfused at 2 mL/min with (in mM) 128 NaCl, 26.2 NaHCO_3_, 2.5 KCl, 1 NaH_2_PO_4_, 1.5 CaCl_2_, 1.5 MgCl_2_, and 11 glucose, maintained at 30.0°C ± 0.5°C. All solutions were equilibrated and maintained with carbogen gas (95% O2/5% CO2; v/v) throughout. Synaptic transmission was routinely blocked by including the following antagonists: 10 μM CNQX (cat #: 1045, Tocris, Minneapolis, MN) AMPA/kainate receptor antagonist, 50 μM d-AP5 (cat #: 0106, Tocris, Minneapolis, MN) NMDA receptor antagonist, and 5 μM SR 95531 hydrobromide (cat #: 1262, Tocris, Minneapolis, MN) GABA_A_ receptor antagonist. Recordings were initiated 15 min after application of synaptic blockers to allow for full pharmacological effect.

### Electrical stimulation hardware

To perform stimulation, a waveform generator (EDU33210A Keysight, Santa Rosa, CA) was used to drive independent current sources (DS4 Digitimer North America, Fort Lauderdale FL). Current was applied to the slice via platinum–iridium electrodes (125 μm diameter, 2 mm length; FHC, USA) for *ex vivo* experiments and via stainless-steel skull screws (E363/96/1.6/SPC, P1 Technologies) for *in vivo* experiment.

### TI stimulation *ex vivo*

Premodulated temporal interference (pmTI) stimulation was applied using sinusoidal waveforms that were generated in advance using Python to produce predefined temporal interference (TI) indices. The TI index was calculated as (Env_max_ − Env_min_)/Env_max_ × 100, where Env_max_ and Env_min_ represent the maximum and minimum amplitudes of the waveform envelope, respectively. The TI index of the waveform was set to 0, 50, or 100.

Stimulation was delivered to cortical layer 2-3 using a bipolar electrode with an inter-electrode spacing of 0.5 mm. Impedance was 7 MΩ at 1 kHz and 1.7 MΩ at 10 kHz. For the 1-s stimulation experiment, premodulated stimulation waveforms generated from 2, 5, and 10 kHz sine waves with a beat frequency of 1 Hz were tested. For each carrier frequency, one stimulation set consisted of sequential application of TI indices AM_0_, AM_50_ and AM_100_ for 1 s with 20-s intervals. This protocol was repeated across different stimulation amplitudes spanning from ±500 μA to ±6 mA, similar to previous studies ^27,70^. In our hands, slice experiments required higher currents to achieve neuronal activation compared with those used in *in vivo* experiments, most likely due to shunting of currents as the electrodes were submerged in electrolytic solution. The nominal electric-field magnitude was estimated to be 60–204 V/cm across the stimulation conditions based on the current, the impedance and the spacing of the electrodes. In a separate experiment, premodulated waveforms generated from a 2-kHz sine wave with a beat frequency of 1 Hz were applied for 10 s for each TI index condition (800–1200 μA; TI indices of 0, 50, and 100%).

For conventional TI stimulation *ex vivo*, two pairs of electrodes were used to deliver currents to generate an interference field within a cortical slice (Supplementary. Fig. 3). Each electrode pair with the 1-mm spacing was placed parallel to the cortical layers, and two pairs were separated by 1.3 mm (Fig. 4a). TI stimulation was generated by delivering two sine-wave currents at 2000 Hz and 2001 Hz through separate electrode pairs, producing a beat frequency of 1 Hz. Stimulation was applied for 5 s with 0.5-s ramp-up and 0.5-s ramp-down periods at amplitudes of ±2–4 mA. As control conditions, sine-wave currents at 2000 Hz were delivered through the same electrode configuration with a 0° phase offset.

### Surgery

Stereotaxic surgery was performed on adult mice (≥110 days old) that had previously received ICV injections of rAAV when they were neonates to prepare them for chronic intravital two-photon calcium imaging of cortical neurons. The surgical protocol was adapted from established procedures for cranial window implantation^71,72^. To enable TI stimulation, bilateral burr hole craniotomies (0.5 mm diameter) were first made anterior and posterior to the planned headplate location at anterior ([AP: +5.5 mm, ML: ±1.0 mm]) and posterior ([AP: −5.5 to −6.5 mm, ML: −1.5 to 0 mm and −3.0 to −4.0 mm]) locations relative to bregma. Stainless-steel skull screws with pre-soldered gold pin attachments (E363/96/1.6/SPC, P1 Technologies) were inserted into the burr holes and fixed to the skull using dental acrylic (C&B Metabond, Parkell). A 3-mm craniotomy was then performed over the anterior left hemisphere to provide optical access to the cortex (AP: −0.3 mm, ML: –1.75 mm). To monitor stimulation artifacts during *in vivo* experiments, an epidural electrode (polyamide insulated 125 mm tungsten wire with exposed tip; P1 Technologies) was inserted at the posterior edge of the craniotomy. A reference stainless screw electrode was also placed contralaterally, superficial to the cerebellum (AP: −5.5 to −6.0 mm, ML: +2.75 to +3.75 mm). A concentric cranial window (inner diameter, 3 mm; outer diameter, 5 mm; D263 #1 thickness coverglass; Warner Instruments) was inserted to seal the craniotomy and secured to the skull using dental acrylic. A stainless-steel headplate (Neurotar) was positioned around the window and affixed to the skull using dental acrylic. Finally, the skin was sealed to the headcap using tissue adhesive (Vetbond, 3M).

### TI stimulation *in vivo*

Skull-screw electrodes were used to deliver sine-wave currents that generated an interference field in the brain. In anesthetized mice, TI stimulation was generated by delivering sine-wave currents at 2000 Hz between the anterior and posterior electrodes on the left side and 2001 or 2000.5-Hz between the anterior and posterior electrodes on the right side. Stimulation was applied for 10-s with 0.5-s ramp-up and 0.5-s ramp-down periods at ±250 –1000 μA. For controls, carrier stimulation at 2000 Hz was delivered through the same electrode configuration. A similar protocol was followed in awake mice, except that stimulation was applied for 30-s with 0.5-s ramp-up and 0.5-s ramp-down periods at current amplitudes of ±50 or ±200 μA. The corresponding nominal electric-field magnitude was estimated to be 2 and 7 V/m for ±50 and ±200 μA, respectively, based on the inter-electrode impedance (∼400 Ω) and the inter-electrode distance (∼12 mm). The ±1 mA stimulation used in the anesthetized mice corresponded to ∼33 V/m. For prolonged stimulation experiments, TI stimulation was generated by delivering sine-wave currents at 2000-Hz on the left side and 2001-Hz on the right side. Stimulation was delivered at ±500 μA in 10-s epochs interleaved with 10-s off periods for a total duration of 20 min, with 0.5-s ramp-up and ramp-down periods for each stimulation epoch. EEG recordings were obtained with an Intan system (Intan, Los Angeles, USA) during stimulation.

### Two-photon imaging

For both *ex vivo* and *in vivo* imaging, resonant galvanometer scanning (30 Hz; 512 x 512 pixels) was performed through a 3-mm working distance 16x water-immersion objective (0.80 numerical aperture, Nikon) using a two-photon microscope (HyperScope, Scientifica) equipped with a pulsed tunable infrared laser system (InSight X3, Spectra-Physics) and controller software (ScanImage, Vidrio Technologies). jGCaMP8s and JEDI-2P were excited using a 925-nm light and fluorescence emission through a dichroic mirror and a band-pass filter (ET525/50m-2p, Chroma) was detected by a GaAsP photomultiplier tube.

*In vivo* imaging was performed either awake or under general anesthesia (2% [v/v] isoflurane balanced in oxygen; flow rate: 1 L/min) at least two weeks following surgery. An awake mouse was head-fixed onto a floating platform (Mobile Home Cage, Neurotar). For imaging under anesthesia, the mouse was head-fixed under the objective with their snout positioned next to a nose cone for gas anesthetic delivery and a heating pad was placed under the mouse to maintain the body temperature during the imaging sessions.

Imaging sessions lasted for 30-90s and consisted of 10-30s of baseline fluorescence imaging followed by 3-30s of stimulation and then 10-30s of a washout period. Recording sessions were separated by at least 1 minute. For prolonged stimulation experiments, calcium imaging was performed continuously for 20 minutes.

### Calcium imaging analyses

Motion registration, region-of-interest (ROI) detection, and extraction of somatic and neuropil fluorescence signals were performed using Suite2P with integrated Cellpose^73,74^. Raw fluorescence traces were corrected for background by subtracting the global minimum of the neuropil signal within each recording. To reduce contamination from out-of-focus fluorescence, the somatic signal was corrected by subtracting 70% of the surrounding neuropil signal. The corrected fluorescence traces were then normalized to ΔF/F₀, where F₀ was defined as the mean fluorescence during the baseline period prior to stimulation onset. For 20-minute recordings, ΔF/F₀ was calculated separately for each 1-min segment using the first 10-s as F₀ to account for photobleaching and baseline drift.

For each cell, stimulus-evoked responses were defined as the peak ΔF/F₀ within predefined stimulation windows. Baseline variability was estimated from the baseline period using a robust standard deviation calculated as 1.4826 × the median absolute deviation (MAD). Responses were classified as significant when they exceeded both the robust z-score threshold (z > 4) and the absolute ΔF/F₀ threshold (ΔF/F₀ > 0.8). Using these criteria, approximately 70% of neurons in the slice experiments and 50% of neurons in the *in vivo* experiments were classified as responsive.

Response thresholds were defined for each neuron as the lowest stimulation amplitude that elicited a stimulus-evoked response under each stimulation condition (AM_0_, AM_50_ and AM_100_). Threshold values were extracted across multiple carrier frequencies (2 kHz, 5 kHz, and 10 kHz). For each neuron, thresholds were included only when responses were observed across all three conditions to ensure consistent comparisons.

Latency following stimulus onset was quantified from 5-Hz low-pass filtered ΔF/F₀ traces. Response onset was defined as the first post-stimulus time point at which the signal exceeded a threshold of the baseline mean plus 3 standard deviations for at least 3 consecutive imaging frames. Latency was measured relative to the onset of a stimulus. For latency analysis, neurons that responded to all stimulation conditions at the same stimulation amplitude were included. For each neuron, latency was quantified at the lowest stimulation amplitude at which this criterion was met.

Powe spectral density (PSD) was estimated for each responsive neuron from ΔF/F₀ traces using Welch’s method (GCaMP8s: 256 samples, 128-sample overlap, nfft = 4096, JEDI2P: window length = 60 samples, no overlap, nfft = 256). PSD was calculated from unfiltered traces within the stimulation window after subtracting the mean of each trace segment. Spectra were restricted to 0.1–5 Hz and normalized such that the total power within this range summed to 1. 1-Hz band power was defined as the normalized power within the 0.8–1.2 Hz band. 0.5-Hz band power was defined as the normalized power within the 0.4–0.6 Hz band.

Event counts were quantified from 5-Hz low-pass filtered calcium signals expressed as ΔF/F_0_. For each ROI and trial, baseline activity level was measured during the first 30-s period before stimulation. A detection threshold was calculated as the baseline median plus zThresh × (1.4826 × MAD), where MAD was the median absolute deviation of the baseline signal. An event was counted when the signal exceeded both this threshold and an absolute ΔF/F_0_ threshold of 0.8 for at least 3 consecutive frames. Event onset was assigned to the first rising point preceding each suprathreshold epoch. ROIs were classified as responsive if they exhibited at least one detected event during the recording period of 90 s, with approximately 60% of ROIs meeting this criterion. Event counts per neuron were then compared across before (0-30s), during (30-60s), and after the stimulation (60-90s).

### Voltage imaging analyses

GEVI fluorescence traces were grouped by slice, and photobleaching was corrected using a baseline-derived single-exponential fit computed separately for each slice. ΔF/F_0_ was then calculated from the corrected fluorescence using the mean signal during the first 10 s of the recording, excluding the first 5 frames, as F_0_. A 5-Hz low-pass Butterworth filter was applied. Mean GEVI ΔF/F_0_ was quantified for each trial by averaging the fluorescence signal during the 3-s stimulation window.

### Statistics

All statistical analyses were performed in MATLAB. Linear mixed-effects models (LMMs) were used for calcium– and voltage-imaging datasets to account for the hierarchical structure of the data. Fixed effects included stimulation condition, carrier frequency, and their interaction, with random intercepts specified according to the structure of each dataset. For slice calcium imaging data, neurons were treated as observations nested within slices and mice. For *in vivo* data, neurons were treated as observations nested within trials and mice. For slice experiments using the voltage indicator, slices were treated as observations nested within trials and mice. For slice imaging with conventional TI dataset (Fig. 4), spatial analyses were performed by binning neurons according to their distance from the interference centerline in 39.6-µm increments. Responsive cell counts were analyzed using a Poisson generalized LMM, whereas the 1-Hz synchronization index was analyzed using LMM. Stimulation condition, distance from the field center, and their interaction were included. For c-Fos experiments, statistical significance was assessed using one-way ANOVA followed by pairwise *t*-tests with Holm–Bonferroni correction. Statistical significance was defined as *p* < 0.05.

### Computational models

We performed simulations in Python using the NEURON44 simulation environment on three neuronal models. These included a previously published biophysical layer 2/3 cortical pyramidal neuron model^26,36,45,46^ and a reduced single-compartment model containing the ion channel composition of axons derived from the same parent model. The published pyramidal neuron model was implemented in NEURON using the Markram/Aberra template and mechanism files, with extracellular stimulation applied through NEURON’s extracellular mechanism. The reduced single-compartment model preserved the principal sodium and potassium conductances and their kinetics in the axon while omitting the full morphology and spatial extracellular field coupling, allowing us to isolate the contribution of axonal membrane dynamics. In uniform-field simulations, stimulation was applied to the published model as a spatially uniform extracellular electric field. For the reduced single-compartment model, the corresponding temporal waveform was delivered as direct current injection.

For conventional TI simulations with the multicompartment model, extracellular potentials were calculated analytically for a four-bar electrode configuration in a homogeneous, isotropic volume conductor with an electrical conductivity of 0.6 S/m. The configuration consisted of four finite bar electrodes, with their near endpoints at x = −10 or +10 mm and y = −10 or +10 mm. For each neuronal segment, the local extracellular potential waveform was calculated from the contributions of both electrode pairs and their time-varying carrier currents at the segment’s actual three-dimensional location and was assigned as the segment-specific extracellular potential.

### Stimulation and c-fos immunohistochemistry

To minimize background c-Fos expression associated with environmental exposure and handling, mice were transferred from the animal facility to the experimental procedure room in covered home cages and anesthetized immediately upon arrival. Mice were kept anesthetized (2-3% [v/v] isoflurane balanced in oxygen; flow rate: 1 L/min) on a heating pad throughout the surgical procedure and stimulation.

Stimulation electrode pairs were custom-fabricated by soldering two spring-loaded gold-plated pins (cat #: 575-906215, Mill-Max, Oyster Bay, NY) to stainless steel wires fixed 1 mm apart using plastic connectors and glue. Following exposure of the skull, small indentations were made in the skull using a hand drill at four locations (AP: −1.94 mm; ML: −4.3, −3.3, −0.78, and +0.22 mm) to facilitate placement of the stimulation electrodes^50,51^. Two pairs of stimulation electrodes were positioned at these sites and secured on the skull using stereotaxic arms. The skull was progressively thinned for each electrode site until an impedance of 1 MΩ was achieved. In the TI stimulation condition, 2,000 Hz and 2,010 Hz currents were delivered through the two stimulation electrode pairs at ±500 μA in 10-s stimulation epochs interleaved with 10-s off periods for a total duration of 20 min. Each stimulation epoch included a 0.5-s ramp-up and a 0.5-s ramp-down. The carrier only stimulation condition underwent the same procedures and received 2,000 Hz currents in phase through both stimulation electrode pairs at the same current amplitude and using the same stimulation protocol. The sham control underwent identical anesthesia and surgical procedures, including electrode placement, but received no electrical stimulation. In separate mice not used for c-Fos quantification, the presence of AM in the hippocampus (AP: –2.7; ML: –2.04; DV: –1.3) was verified by inserting a recording electrode. The electrode was inserted at a 20° angle in the anteroposterior direction. Intracranial recordings were obtained with an Intan system (Intan Technologies, Los Angeles, CA, USA) using stainless-steel recording electrodes (MS303/3-B/SPC, 0.005-inch wire diameter, Protech International) during TI stimulation with ±400 µA currents at 2000 Hz and 2005 Hz.

Ninety minutes post end of stimulation, mice were sacrificed by IP injection of pentobarbital/phenytoin solution (0.5 mL, Virbac Euthasol, 1:39 dilution of stock solution in saline) and transcardiac perfusion with PBS and 4% (w/v) paraformaldehyde (PFA). Brains dissected from the skull and after at least 24 hours in PFA were moved to 30% (w/v) sucrose solution until sinking for cryoprotection. Brains were then sectioned coronally on a cryostat at 50-µm thickness, collected in 8 series, and retained as floating sections. Brains from one series were permeabilized in 3% (v/v) H_2_O_2_ and 0.1% (v/v) Triton X-100 in PBS for 30 minutes, rinsed 3 times in PBS for 10 minutes and blocked in 4% (v/v) normal donkey serum in PBS for 30 minutes. They were then incubated in primary antibody (Rabbit anti-c-Fos, 1:500 dilution, Abcam cat #: AB222699 1 ug/ml) at 4°C overnight, rinsed, and incubated in a biotinylated donkey anti-rabbit secondary antibody (Biotin-SP-conjugated donkey anti-Rabbit IgG, Jackson ImmunoResearch Laboratories, cat #: 711-065-152, 4 µg/mL) for an hour at room temperature, rinsed, and incubated with an avidin-bound detection enzyme (Vectastain ABC-HRP Kit #: PK-4001) at 4°C overnight. Sections were then rinsed, and staining was completed with addition of the chromogen (3,3’-diaminobenzidine tetrachloride) for 12 minutes in the presence of H_2_O_2_. After the final rinse, the sections were mounted on glass slides, dehydrated, and coverslipped. All slides were digitally imaged using a slide scanner (Leica Aperio AT2).

Quantification of c-Fos-positive cells was performed using open-source software (QuPath v0.6.0). All analyses were performed by an investigator blinded to the experimental conditions. Counts were performed at antero-posterior levels corresponding to the region under the stimulation electrodes. Regions of interest (ROIs) within the cortex, CA1 and DG were manually annotated. Color deconvolution was applied to estimate stain vectors for DAB. Automated cell detection was executed using the thresholds for optical density set to distinguish positive from negative cells. Data including cell counts within the defined ROIs and the size of ROIs were exported to calculate density.

**Supplementary Fig. 1.**
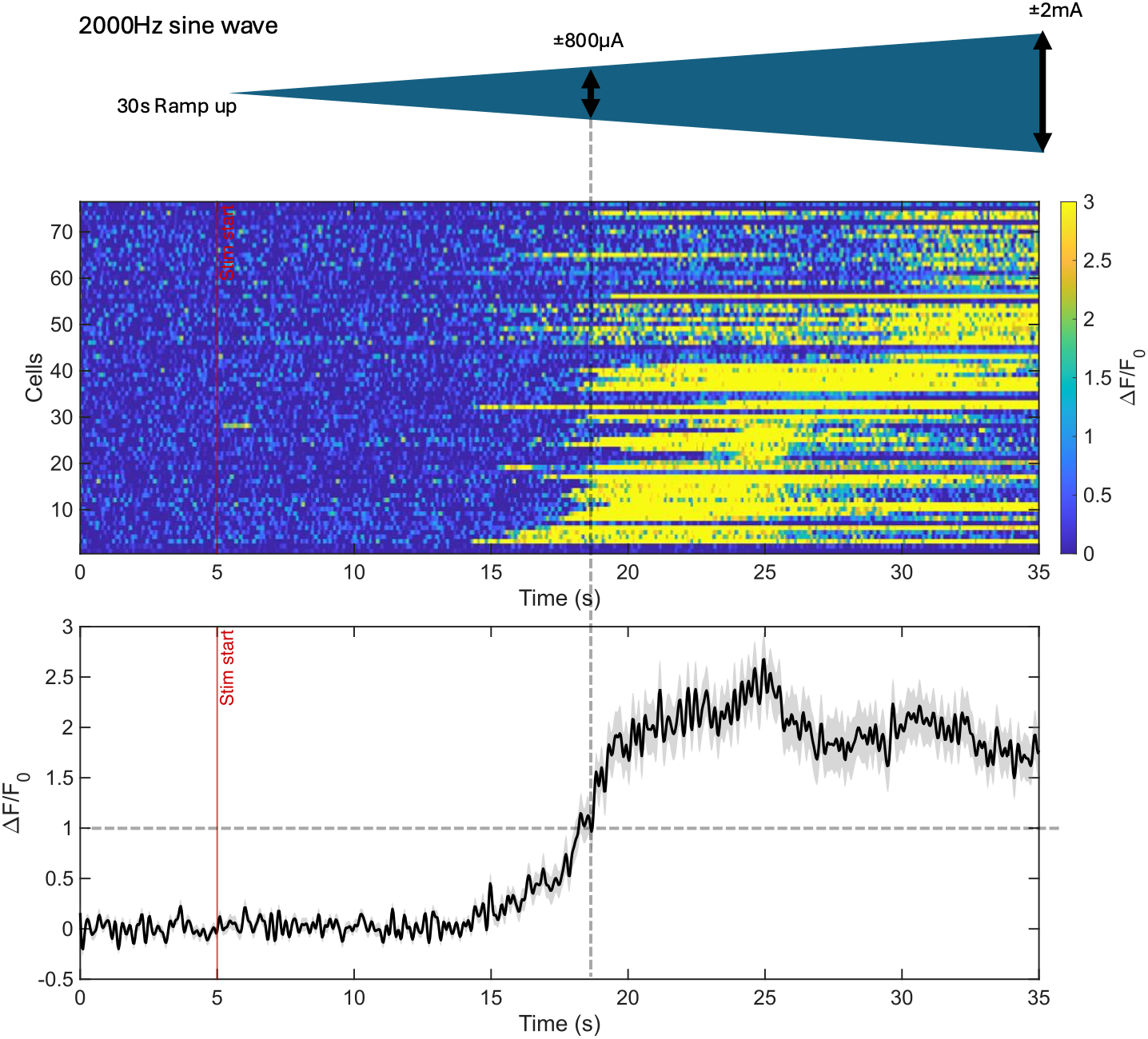
Neurons showed responses during long-ramp-up stimulation regardless of the gradually increasing stimulus amplitude. After a 5-s baseline, 2000Hz sine wave stimulation amplitude was linearly ramped for 30-s. Top, schematic of the long-ramp-up stimulation waveform, increasing linearly from 0 to ±2 mA over 30-s. Middle, ΔF/F_0_ heatmap for cells (n = 76). Color indicates ΔF/F_0_. Bottom, population mean ΔF/F_0_ across all cells (black); shaded area indicates SEM.

**Supplementary Fig. 2.**
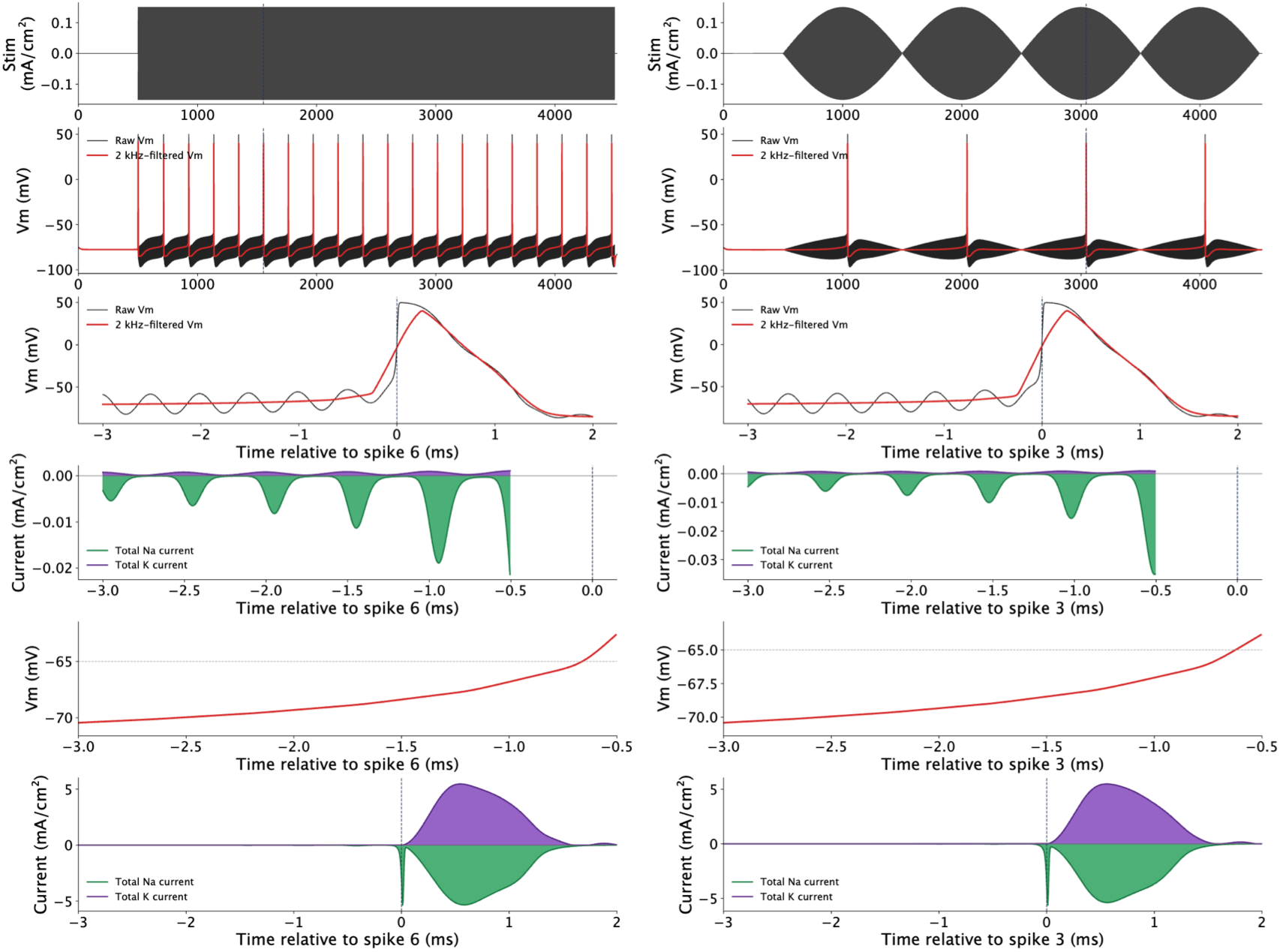
Simulated responses of Reduced single-compartment axonal model to high-frequency stimulation. Left, reduced single-compartment axonal model under unmodulated carrier stimulation. Right, reduced single-compartment axonal model under TI stimulation. From top to bottom, traces show: (1) the applied stimulation waveform; (2) membrane potential over the full stimulation period, with the 2-kHz-filtered membrane potential shown in red; (3) membrane potential immediately preceding and during spike initiation; (4) sodium and potassium currents during the corresponding pre-spike period; (5) the filtered membrane potential during the subthreshold period preceding spike initiation; and (6) the corresponding filtered sodium and potassium currents. Green indicates inward sodium current and purple indicates outward potassium current.

**Supplementary Fig. 3.**
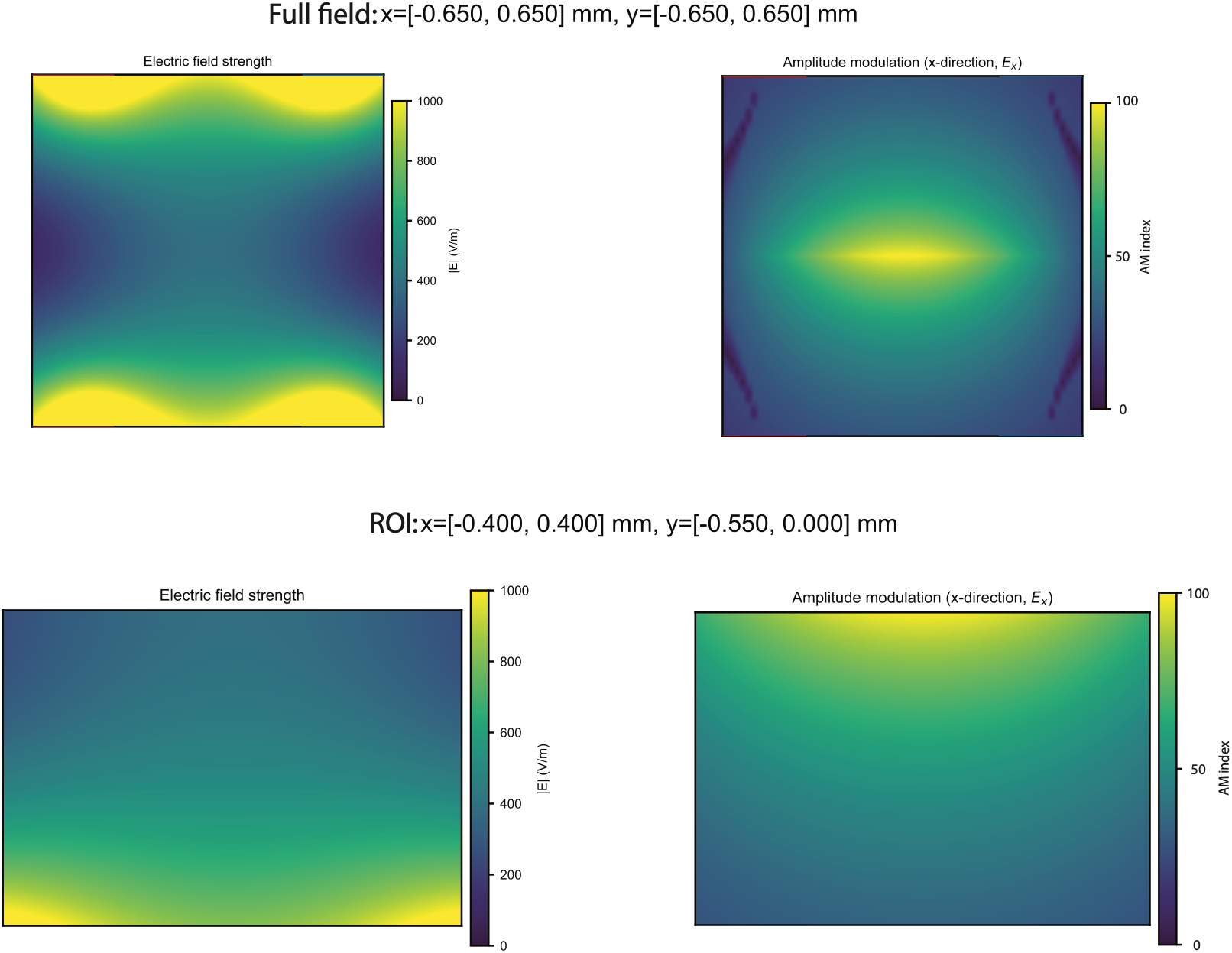
Simulated electric field and AM index for the conventional TI electrode configuration *ex vivo*. Left panels show the electric-field magnitude (|E|), and right panels show the normalized AM index of the x-component of the electric field (Ex). The top row shows the full field distribution (x = −0.65 to 0.65 mm, y = −0.65 to 0.65 mm), whereas the bottom row shows the region of interest (ROI; x = −0.40 to 0.40 mm, y = −0.55 to 0 mm) used for spatial analysis. The two electrode pairs were positioned at y = −0.65 and 0.65 mm, respectively, with the electrodes within each pair located at x = −0.5 and 0.5 mm. Simulations were performed with a 1-mm intra-pair electrode spacing, a 1.3-mm inter-pair distance, a 2-mm exposed electrode length, a uniform conductivity of 0.6 S/m, and a stimulation amplitude of 4 mA per electrode.

**Supplementary Fig. 4.**
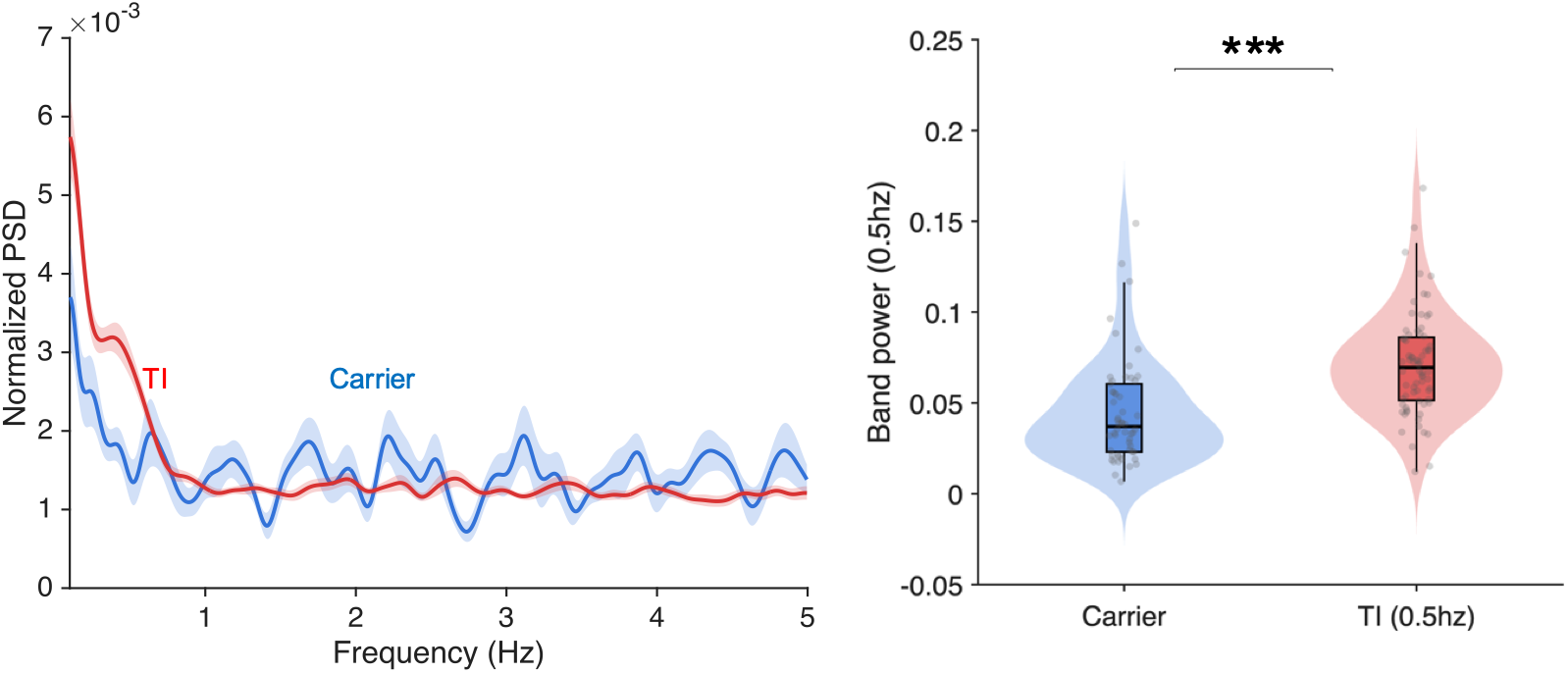
0.5-Hz TI stimulation enhances neuronal entrainment at the beat frequency relative to carrier stimulation. Comparison of normalized PSD power in responsive neurons during carrier and TI (0.5-Hz) stimulation. Left, normalized PSD spectra from 0.1 to 5-Hz for carrier (2-kHz; blue) and TI (0.5-Hz; red) stimulation. Thick traces indicate the mean and shaded areas indicate SEM across responsive neurons; the dashed line marks 0.5-Hz. Right, quantification of normalized 0.5-Hz band power (0.4–0.6-Hz). Violin plots show the distribution for each condition; box plots indicate the median and interquartile range, and dots indicate individual cells. LMM, p < 0.05; n = 72 cells for TI and n = 54 cells for carrier from 3 mice.

**Supplementary Fig. 5.**
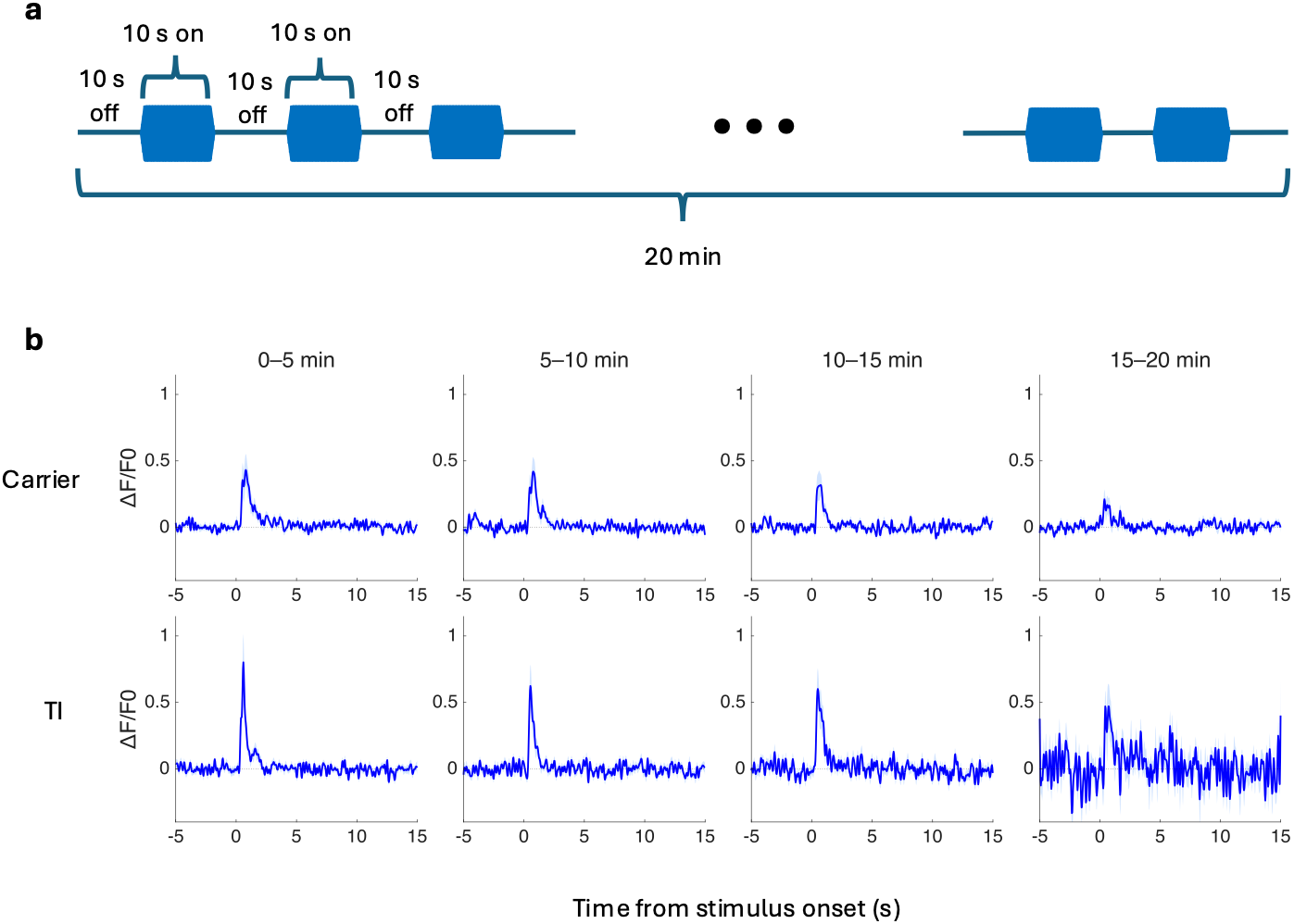
Sustained calcium responses during 20-min carrier and TI stimulation. **a**, stimulation paradigm: 10-s stimulation ON and 10-s OFF cycles were repeated for 20min. Carrier (2-kHz) or TI (2000 and 2010-Hz; 10-Hz beat frequency) was kept constant throughout each recording. b, mean ΔF/F_0_ traces of all cells after stimulus onset, shown separately for four 5-min epochs (0– 5, 5–10, 10–15, and 15–20-min). The upper row shows responses during carrier stimulation, and the lower row shows responses during TI stimulation. Data are from one mouse (28 cells).

**Supplementary Fig. 6.**
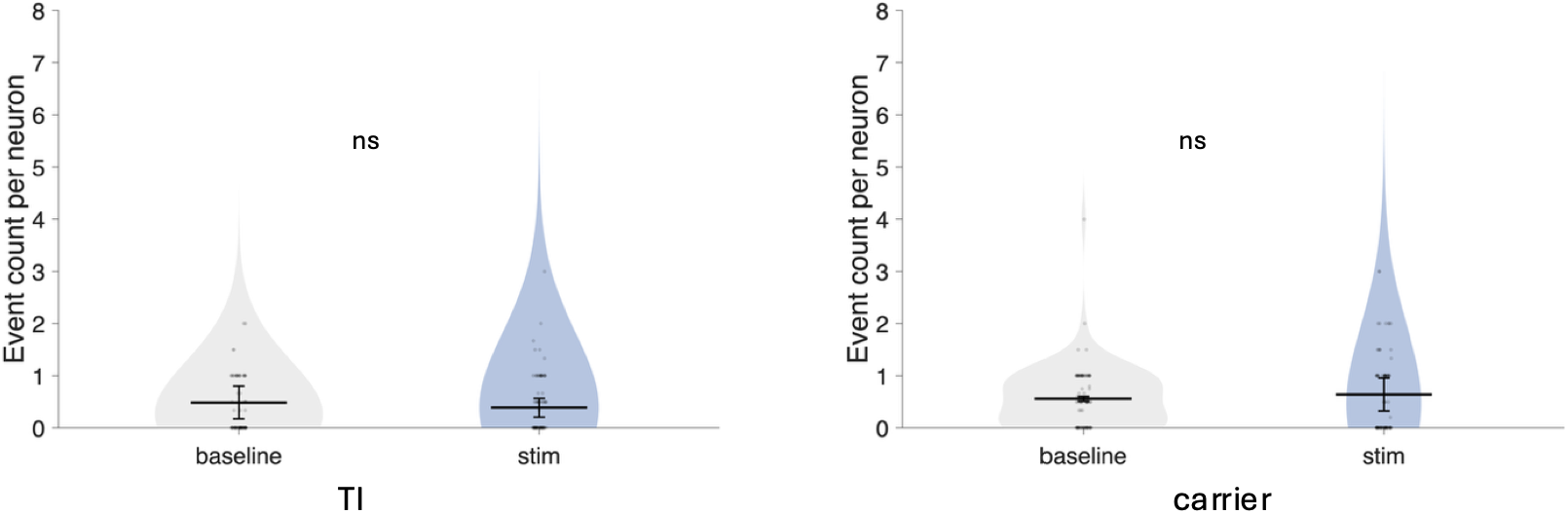
250 uA did not induce a significant direct response. Event count per neuron during baseline (0-10 s) and stimulation (10-20 s) at 250 uA. Responsive ROIs were defined as neurons showing at least one detected event within 0-30 s in the 200 uA condition. Counts were extracted from traces and compared using a linear mixed-effects model. No significant increase in event count during stimulation was detected in either TI (n = 78 neurons, 3 mice, p = 0.322) or carrier (n = 88 neurons, 3 mice, p = 0.773).

**Supplementary Fig. 7.**
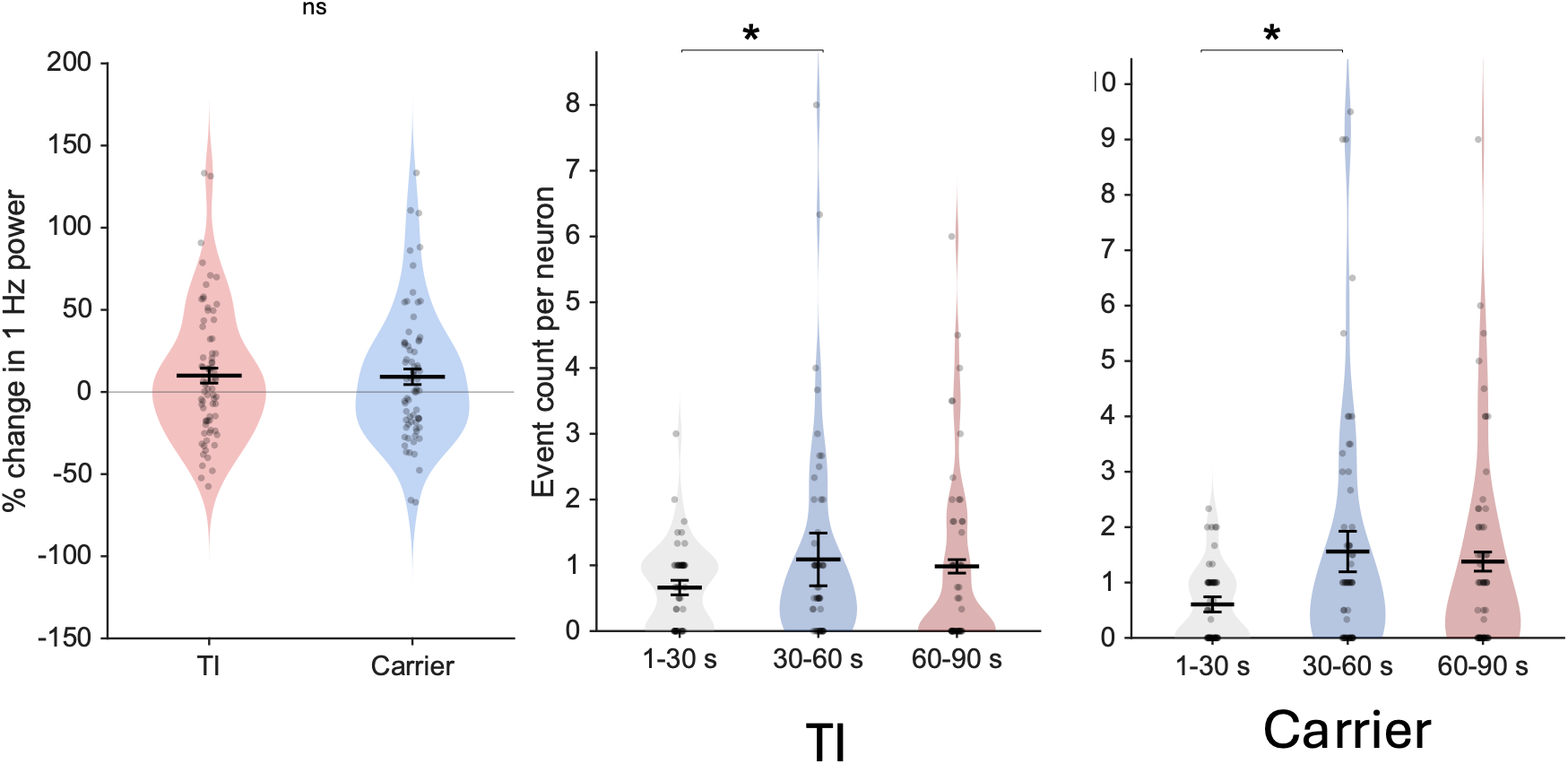
5kHz carrier stimulation excited neuronal activity without a differential change in 1-Hz power. Left. Percent change in normalized 1-Hz power from the baseline period (0–30 s) to the stimulation period (30–60 s) in responsive neurons during TI and carrier (5 kHz) stimulation. Each dot represents one neuron; violins show distributions, and black bars indicate mean ± SEM. No significant difference was detected between TI and carrier stimulation (ns). Middle and Right. Event counts per responsive neuron during the 1–30 s, 30–60 s, and 60–90 s epochs for TI (Middle). and carrier (5 kHz) stimulation (Right), respectively. Each dot represents one neuron; violins show distributions, and black bars indicate mean ± SEM. Statistical comparisons were performed using LMM. *p* < 0.05. n = 74 cells for TI, n = 71 cells for carrier from 3 mice

## Acknowledgments

We thank Anna Blomerth and Broderick Barnes for their assistance with histology. This work was supported by funding from the NIH [R01NS13873302 (CAG, KB), S10OD021773 (KB)], The Emory NeuroNetworks Initiative (CAG, NL). and Japan Student Services Organization (RY).

## Author contributions

Conceptualization: RY, KB, CAG, MAS; Methodology: RY, KB, CAG, MAS; Software: RY, MAS; Validation: RY, KB; Formal analysis: RY, KB, CAG, MAS; Investigation: RY, MAS, KB, CAG; Resources: CAG, KB; Data curation: RY, KB, CAG; Writing—original draft: RY; Writing—review and editing: RY, KB, CAG, MAS, TE, EA, NL; Visualization: RY, KB, CAG; Supervision: KB, CAG; Project administration: RY; Funding acquisition: CAG, KB, EA

## Data and materials availability

Code and summary data needed to reproduce the analyses and simulations presented in this study will be publicly archived at DANDI [repository pending, to be updated prior to publication].

## BIBLIOGRAPHY AND REFERENCES CITED

1 Lozano, A. M., et al. Deep brain stimulation: current challenges and future directions. Nat Rev Neurol 15, 148–160 (2019). 10.1038/s41582-018-0128-2

2 Krauss, J. K., et al. Technology of deep brain stimulation: current status and future directions. Nat Rev Neurol 17, 75–87 (2021). 10.1038/s41582-020-00426-z

3 Deshpande, N., Hadi, M. & Ali, R. Healthcare disparities in deep brain stimulation access and utilization: a systematic review. J Neurosurg 140, 1137–1147 (2024). 10.3171/2023.7.JNS23277

4 Fenoy, A. J. & Simpson, R. K., Jr. Risks of common complications in deep brain stimulation surgery: management and avoidance. J Neurosurg 120, 132–139 (2014). 10.3171/2013.10.JNS131225

5 Fertonani, A. & Miniussi, C. Transcranial Electrical Stimulation: What We Know and Do Not Know About Mechanisms. Neuroscientist 23, 109–123 (2017). 10.1177/1073858416631966

6 Hallett, M. Transcranial magnetic stimulation: a primer. Neuron 55, 187–199 (2007). 10.1016/j.neuron.2007.06.026

7 Schulz, R., Gerloff, C. & Hummel, F. C. Non-invasive brain stimulation in neurological diseases. Neuropharmacology 64, 579–587 (2013). 10.1016/j.neuropharm.2012.05.016

8 Voroslakos, M., et al. Direct effects of transcranial electric stimulation on brain circuits in rats and humans. Nat Commun 9, 483 (2018). 10.1038/s41467-018-02928-3

9 Deng, Z. D., Lisanby, S. H. & Peterchev, A. V. Electric field depth-focality tradeoff in transcranial magnetic stimulation: simulation comparison of 50 coil designs. Brain Stimul 6, 1–13 (2013). 10.1016/j.brs.2012.02.005

10 Grossman, N., et al. Noninvasive Deep Brain Stimulation via Temporally Interfering Electric Fields. Cell 169, 1029–1041 e1016 (2017). 10.1016/j.cell.2017.05.024

11 Hutcheon, B. & Yarom, Y. Resonance, oscillation and the intrinsic frequency preferences of neurons. Trends Neurosci 23, 216–222 (2000). 10.1016/s0166-2236(00)01547-2

12 Demchenko, I., et al. Human applications of transcranial temporal interference stimulation: A systematic review. Brain Stimul 18, 2054–2066 (2025). 10.1016/j.brs.2025.10.023

13 Liu, R., et al. Temporal interference stimulation targets deep primate brain. Neuroimage 291, 120581 (2024). 10.1016/j.neuroimage.2024.120581

14 Vieira, P. G., Krause, M. R. & Pack, C. C. Temporal interference stimulation disrupts spike timing in the primate brain. Nat Commun 15, 4558 (2024). 10.1038/s41467-024-48962-2

15 Beanato, E., et al. Noninvasive modulation of the hippocampal-entorhinal complex during spatial navigation in humans. Sci Adv 10, eado4103 (2024). 10.1126/sciadv.ado4103

16 Violante, I. R., et al. Non-invasive temporal interference electrical stimulation of the human hippocampus. Nat Neurosci 26, 1994–2004 (2023). 10.1038/s41593-023-01456-8

17 Wessel, M. J., et al. Noninvasive theta-burst stimulation of the human striatum enhances striatal activity and motor skill learning. Nat Neurosci 26, 2005–2016 (2023). 10.1038/s41593-023-01457-7

18 Vassiliadis, P., et al. Non-invasive stimulation of the human striatum disrupts reinforcement learning of motor skills. Nat Hum Behav 8, 1581–1598 (2024). 10.1038/s41562-024-01901-z

19 Ryan, J. S., et al. Ventral striatum temporal interference brain stimulation enhances the reward-positivity event-related potential and reduces anxiety. Neuroimage 338, 122101 (2026). 10.1016/j.neuroimage.2026.122101

20 Missey, F., et al. Non-invasive temporal interference stimulation of the hippocampus suppresses epileptic biomarkers in patients with Epilepsy: biophysical differences between kilohertz and amplitude modulated stimulation. Brain Stimul 19, 102981 (2026). 10.1016/j.brs.2025.11.008

21 Yang, C., et al. Transcranial Temporal Interference Stimulation of the Right Globus Pallidus in Parkinson’s Disease. Mov Disord 40, 1061–1069 (2025). 10.1002/mds.29967

22 Lamos, M., et al. Noninvasive Temporal Interference Stimulation of the Subthalamic Nucleus in Parkinson’s Disease Reduces Beta Activity. Mov Disord 40, 1051–1060 (2025). 10.1002/mds.30134

23 Fani, N. & Treadway, M. T. Potential applications of temporal interference deep brain stimulation for the treatment of transdiagnostic conditions in psychiatry. Neuropsychopharmacology 49, 305–306 (2024). 10.1038/s41386-023-01682-5

24 Modak, P., et al. Temporal interference electrical neurostimulation at 20 Hz beat frequency leads to increased fMRI BOLD activation in orbitofrontal cortex in humans. Brain Stimul 17, 867–875 (2024). 10.1016/j.brs.2024.07.014

25 Savvateev, I., et al. Multipair phase-modulated temporal interference electrical stimulation combined with fMRI. Cell Syst, 101610 (2026). 10.1016/j.cels.2026.101610

26 Caldas-Martinez, S., et al. Cell-specific effects of temporal interference stimulation on cortical function. Commun Biol 7, 1076 (2024). 10.1038/s42003-024-06728-y

27 Semenov, I., Kim, V., Silkuniene, G. & Pakhomov, A. G. Excitation and polarization of isolated neurons by high-frequency sine waves for temporal interference stimulation. Cell Rep Phys Sci 6 (2025). 10.1016/j.xcrp.2025.102660

28 Luff, C. E., Dzialecka, P., Acerbo, E., Williamson, A. & Grossman, N. Pulse-width modulated temporal interference (PWM-TI) brain stimulation. Brain Stimul 17, 92–103 (2024). 10.1016/j.brs.2023.12.010

29 Luff, C. E., et al. The neuron mixer and its impact on human brain dynamics. Cell Rep 43, 114274 (2024). 10.1016/j.celrep.2024.114274

30 Esmaeilpour, Z., Kronberg, G., Reato, D., Parra, L. C. & Bikson, M. Temporal interference stimulation targets deep brain regions by modulating neural oscillations. Brain Stimul 14, 55–65 (2021). 10.1016/j.brs.2020.11.007

31 Katz, B. Nerve excitation by high-frequency alternating current. J Physiol 96, 202–224 (1939). 10.1113/jphysiol.1939.sp003771

32 Chaieb, L., Antal, A. & Paulus, W. Transcranial alternating current stimulation in the low kHz range increases motor cortex excitability. Restor Neurol Neurosci 29, 167–175 (2011). 10.3233/RNN-2011-0589

33. Ravasio, C. R., et al. Kilohertz electrical stimulation evokes robust cellular responses like conventional frequencies but distinct population dynamics. Commun Biol 8, 19 (2025). 10.1038/s42003-024-07447-0

34 Opancar, A., et al. The same biophysical mechanism is involved in both temporal interference and direct kHz stimulation of peripheral nerves. Nat Commun 16, 9006 (2025). 10.1038/s41467-025-64059-w

35 Mirzakhalili, E., Barra, B., Capogrosso, M. & Lempka, S. F. Biophysics of Temporal Interference Stimulation. Cell Syst 11, 557–572 e555 (2020). 10.1016/j.cels.2020.10.004

36 Wang, B., Aberra, A. S., Grill, W. M. & Peterchev, A. V. Responses of model cortical neurons to temporal interference stimulation and related transcranial alternating current stimulation modalities. J Neural Eng 19 (2023). 10.1088/1741-2552/acab30

37 Labruna, L., et al. Kilohertz transcranial magnetic perturbation (kTMP) as a new non-invasive method to modulate cortical excitability. Elife 13 (2025). 10.7554/eLife.92088

38 Kilgore, K. L. & Bhadra, N. Reversible nerve conduction block using kilohertz frequency alternating current. Neuromodulation 17, 242–254; discussion 254–245 (2014). 10.1111/ner.12100

39 Cassara, A. M., et al. Recommendations for the Safe Application of Temporal Interference Stimulation in the Human Brain Part II: Biophysics, Dosimetry, and Safety Recommendations. Bioelectromagnetics 46, e22536 (2025). 10.1002/bem.22536

40 Howell, B. & McIntyre, C. C. Feasibility of Interferential and Pulsed Transcranial Electrical Stimulation for Neuromodulation at the Human Scale. Neuromodulation 24, 843–853 (2021). 10.1111/ner.13137

41 Rampersad, S., et al. Prospects for transcranial temporal interference stimulation in humans: A computational study. Neuroimage 202, 116124 (2019). 10.1016/j.neuroimage.2019.116124

42 Eggers, T., Acerbo, E., Laxpati, N. & Gutekunst, C. A. Direct delivery of modulated kilohertz frequency waveforms enable simultaneous electrical stimulation and recording with minimal-artifact. J Neural Eng 23 (2026). 10.1088/1741-2552/ae7765

43 Zhang, Y., et al. Fast and sensitive GCaMP calcium indicators for imaging neural populations. Nature 615, 884–891 (2023). 10.1038/s41586-023-05828-9

44 Hines, M. L. & Carnevale, N. T. The NEURON simulation environment. Neural Comput 9, 1179–1209 (1997). 10.1162/neco.1997.9.6.1179

45 Aberra, A. S., Peterchev, A. V. & Grill, W. M. Biophysically realistic neuron models for simulation of cortical stimulation. J Neural Eng 15, 066023 (2018). 10.1088/1741-2552/aadbb1

46 Markram, H., et al. Reconstruction and Simulation of Neocortical Microcircuitry. Cell 163, 456–492 (2015). 10.1016/j.cell.2015.09.029

47 Hou, X., et al. Nanobubble-actuated ultrasound neuromodulation for selectively shaping behavior in mice. Nat Commun 15, 2253 (2024). 10.1038/s41467-024-46461-y

48. Grotemeyer, A., et al. Standardized wireless deep brain stimulation system for mice. NPJ Parkinsons Dis 10, 153 (2024). 10.1038/s41531-024-00767-2

49 Carmona-Barron, V. G., et al. Comparing the effects of transcranial alternating current and temporal interference (tTIS) electric stimulation through whole-brain mapping of c-Fos immunoreactivity. Front Neuroanat 17, 1128193 (2023). 10.3389/fnana.2023.1128193

50 Missey, F., et al. Orientation of Temporal Interference for Non-invasive Deep Brain Stimulation in Epilepsy. Front Neurosci 15, 633988 (2021). 10.3389/fnins.2021.633988

51 Acerbo, E., et al. Focal non-invasive deep-brain stimulation with temporal interference for the suppression of epileptic biomarkers. Front Neurosci 16, 945221 (2022). 10.3389/fnins.2022.945221

52 Liu, Z., et al. Sustained deep-tissue voltage recording using a fast indicator evolved for two-photon microscopy. Cell 185, 3408–3425 e3429 (2022). 10.1016/j.cell.2022.07.013

53 Ackermann, D. M., Bhadra, N., Gerges, M. & Thomas, P. J. Dynamics and sensitivity analysis of high-frequency conduction block. J Neural Eng 8, 065007 (2011). 10.1088/1741-2560/8/6/065007

54 Johnson, L., et al. Dose-dependent effects of transcranial alternating current stimulation on spike timing in awake nonhuman primates. Sci Adv 6 (2020). 10.1126/sciadv.aaz2747

55 Tran, H., Shirinpour, S. & Opitz, A. Effects of transcranial alternating current stimulation on spiking activity in computational models of single neocortical neurons. Neuroimage 250, 118953 (2022). 10.1016/j.neuroimage.2022.118953

56 Krause, M. R., Vieira, P. G., Thivierge, J. P. & Pack, C. C. Brain stimulation competes with ongoing oscillations for control of spike timing in the primate brain. PLoS Biol 20, e3001650 (2022). 10.1371/journal.pbio.3001650

57 de Graaf, T. A., et al. Alpha-band rhythms in visual task performance: phase-locking by rhythmic sensory stimulation. PLoS One 8, e60035 (2013). 10.1371/journal.pone.0060035

58 Alagapan, S., et al. Modulation of Cortical Oscillations by Low-Frequency Direct Cortical Stimulation Is State-Dependent. PLoS Biol 14, e1002424 (2016). 10.1371/journal.pbio.1002424

59 Neuling, T., Rach, S. & Herrmann, C. S. Orchestrating neuronal networks: sustained after-effects of transcranial alternating current stimulation depend upon brain states. Front Hum Neurosci 7, 161 (2013). 10.3389/fnhum.2013.00161

60 Silvanto, J. & Pascual-Leone, A. State-dependency of transcranial magnetic stimulation. Brain Topogr 21, 1–10 (2008). 10.1007/s10548-008-0067-0

61 Logothetis, N. K., Pauls, J., Augath, M., Trinath, T. & Oeltermann, A. Neurophysiological investigation of the basis of the fMRI signal. Nature 412, 150–157 (2001). 10.1038/35084005

62 Goense, J. B. & Logothetis, N. K. Neurophysiology of the BOLD fMRI signal in awake monkeys. Curr Biol 18, 631–640 (2008). 10.1016/j.cub.2008.03.054

63 Bartolo, M. J., et al. Stimulus-induced dissociation of neuronal firing rates and local field potential gamma power and its relationship to the resonance blood oxygen level-dependent signal in macaque primary visual cortex. Eur J Neurosci 34, 1857–1870 (2011). 10.1111/j.1460-9568.2011.07877.x

64 Botzanowski, B., et al. Focal control of non-invasive deep brain stimulation using multipolar temporal interference. Bioelectron Med 11, 7 (2025). 10.1186/s42234-025-00169-6

65 Song, X., Zhao, X., Li, X., Liu, S. & Ming, D. Multi-channel transcranial temporally interfering stimulation (tTIS): application to living mice brain. J Neural Eng 18 (2021). 10.1088/1741-2552/abd2c9

66 Mencarelli, L., et al. Impact of network-targeted multichannel transcranial direct current stimulation on intrinsic and network-to-network functional connectivity. J Neurosci Res 98, 1843–1856 (2020). 10.1002/jnr.24690

67 Wang, J. X., et al. Targeted enhancement of cortical-hippocampal brain networks and associative memory. Science 345, 1054–1057 (2014). 10.1126/science.1252900

68 Fox, M. D., Buckner, R. L., White, M. P., Greicius, M. D. & Pascual-Leone, A. Efficacy of transcranial magnetic stimulation targets for depression is related to intrinsic functional connectivity with the subgenual cingulate. Biol Psychiatry 72, 595–603 (2012). 10.1016/j.biopsych.2012.04.028

69 Kim, J. Y., Grunke, S. D., Levites, Y., Golde, T. E. & Jankowsky, J. L. Intracerebroventricular viral injection of the neonatal mouse brain for persistent and widespread neuronal transduction. J Vis Exp, 51863 (2014). 10.3791/51863

70 Lu, H., Shaner, S., Otte, E., Asplund, M. & Vlachos, A. A microfluidic perspective on conventional in vitro transcranial direct current stimulation methods. J Neurosci Methods 385, 109761 (2023). 10.1016/j.jneumeth.2022.109761

71 Stern, M. A., Cole, E. R., Gross, R. E. & Berglund, K. Seizure event detection using intravital two-photon calcium imaging data. Neurophotonics 11, 024202 (2024). 10.1117/1.NPh.11.2.024202

72 Goldey, G. J., et al. Removable cranial windows for long-term imaging in awake mice. Nat Protoc 9, 2515–2538 (2014). 10.1038/nprot.2014.165

73 Stringer, C., Wang, T., Michaelos, M. & Pachitariu, M. Cellpose: a generalist algorithm for cellular segmentation. Nat Methods 18, 100–106 (2021). 10.1038/s41592-020-01018-x

74 Pachitariu, M. et al. Suite2p: beyond 10,000 neurons with standard two-photon microscopy. bioRxiv, 061507 (2017). 10.1101/061507

